# An antimicrobial peptide expression platform for targeting pathogenic bacterial species

**DOI:** 10.1101/2023.10.09.561505

**Authors:** Jack W. Rutter, Linda Dekker, Chania Clare, Julie A.K. McDonald, Sean P. Nair, Alex J.H. Fedorec, Chris P. Barnes

**Affiliations:** Department of Cell and Developmental Biology, University College London, London, UK; Division of Digestive Diseases, Department of Metabolism, Digestion and Reproduction, Imperial College London, London, UK; Division of Microbial Diseases, UCL Eastman Dental Institute, University College London, London, UK

**Keywords:** engineering biology, antimicrobial peptides, human disease, microbiota, ESKAPE pathogens

## Abstract

Bacteriocins are antimicrobial peptides that are naturally produced by many bacteria. They hold great potential in the fight against antibiotic resistant bacteria, including ESKAPE pathogens. However, they often have low stability *in vivo* and therefore, may not be effective when administered orally. Engineered live biotherapeutic products (eLBPs) that secrete bacteriocins can be created to deliver bacteriocins directly at the site of infection. Here we develop a modular bacteriocin secretion platform that can be used to express and secrete multiple bacteriocins from non-pathogenic *Escherichia coli* host strains. As a proof of concept we create Enterocin A and Enterocin B secreting strains that show strong antimicrobial activity against *Enterococcus faecalis* and *Enterococcus faecium*, and characterise this activity in both solid culture and liquid co-culture. We then develop a Lokta-Volterra model that can be used to capture the interactions of these competitor strains. We show that simultaneous exposure to EntA and EntB can delay the point of *Enterococcus* growth. Our system has the potential to be used as an eLBP to secrete additional bacteriocins for the targeted killing of other pathogenic bacteria.

## 1 Introduction

The microbiota has a profound impact on human health and these communities are implicated in many pathological states. This impact has created growing interest in methods that can manipulate the human host-microbiota system to combat disease(*1* –*4*). One approach is to use engineering biology techniques to construct engineered live biotherapeutic products (eLBPs)(*5*). These eLBPs are typically bacterial strains that have been modified to perform a therapeutic purpose, e.g. the production of a therapeutic molecule. Currently many eLBPs are undergoing clinical trials to assess their efficacy in human patients, with indications for cancer and hyperoxaluria, amongst others(*6*). Another promising application of eLBPs is the production of antimicrobial peptides (AMPs) that can be used to target pathogenic species.

Alongside technologies such as phage therapy(*7*), AMPs have emerged as a potential alternative to traditional antibiotics in the face of concerns over antibiotic resistance(*8*). AMPs are a class of small peptides that display antimicrobial activity against a range of bacteria, fungi, viruses and parasites. Within this study we focus on a subset of AMPs known as bacteriocins(*9*). All major lineages of bacteria are thought to produce at least one bacteriocin(*10*) for self-preservation or to produce a competitive advantage in polymicrobial environments(*11*). Bacteriocins are ribosomally synthesised and generally show potent activity against a narrow spectrum of bacteria closely related to the producing species (although some broad-spectrum bacteriocins do exist)(*12*). This narrow spectrum of activity is a desirable trait for human health applications; as narrow spectrum treatments allow for the targeted removal of pathogenic species, while limiting the disruptive impact the antimicrobial has on the native microbiota.

To date, bacteriocins have been used in a range of biotechnological and industrial applications. However, they have not yet been used extensively in health applications(*13*). Bacteriocins are readily degraded by proteases in the mammalian digestive tract and show reduced stability across different pH values, and it is challenging to ensure sufficient quantities reach the infection site when administered orally(*14*, *15*). Alternative routes, such as intravenous or subcutaneous delivery are also difficult, as bloodstream proteases can reduce bacteriocin activity and injection-site degradation may occur(*13*, *14*). The development of eLBP platforms that can be used to deliver bacteriocins directly at the site of infection have the potential to overcome these challenges. A suitable secretion system is required to ensure that the expressed bacteriocin is able to exit the host cell and target pathogenic bacteria in the environment. It has previously been shown that secretion signals affect the synthesis and secretion of proteins in an unpredictable manner(*16*). Although some studies have applied machine learning techniques to the classification and *de novo* design of secretion peptides(*17*, *18*), it is still a non-trivial task to predict the optimal secretion signal for a specific peptide sequence(*19*).

Bacteriocin producing systems have been previously reported (*20* –*22*). Geldart *et al* designed the “pMPES2” platform, which repurposed the microcin V (mccV) secretion system to express multiple bacteriocins from a single operon under the control of the constitutive ProTeOn+ promoter(*23*, *24*). The authors showed that simultaneously producing multiple bacteriocins was able to slow the development of resistance in the target species. The strain also reduced *E. faecium* 8E9 and *E. faecalis* V583R colonisation in a mouse model of vancomycin-resistant enterococci (VRE) infection. Although the system was successful, it was shown that the operon layout of multiple-bacteriocin producing strains greatly affected the antimicrobial activity of their pMPES2 platform(*24*).

In this study we report the development of a new bacteriocin secretion platform that can be used to target pathogenic species. As a proof of concept we target two *Enterococcus* species as pathogens of interest, *Enterococcus faecalis* and *Enterococcus faecium*. One of the most prevalent enterococci strains found in the human gut(*25*), *E. faecalis* is a grampositive, opportunistic pathogen that is associated with endocarditis, systemic and urinary tract infections(*10*). Furthermore, there is some evidence that *E. faecalis* is involved in the development of colorectal cancer(*25*), although there is strong debate whether it plays a harmful or protective role(*26* –*30*). *E. faecium*, identified as an ESKAPE pathogen by the World Health Organisation, has been shown to exhibit resistance to a range of antibiotics and is the most common causative agent of VRE infections(*31* –*33*). We build on previous works in two main ways. We develop a modular platform based on the CIDAR MoClo assembly standard(*34*), which allows for flexibility in part interchange. Additionally we explore four different secretion signal peptides, MalE, OmpA, PhoA and PM3 (a modified version of the PelB secretion tag(*35*)), to add another tunable parameter for eLBP delivery. We then use co-culture assays and Lotka-Volterra modeling to gain insights into the growth dynamics of these strains.

## 2 Results

### 2.1 Development of the AMP system

We started by designing a modular platform for the expression and secretion of bacteriocins. A summary of this bacteriocin expression platform is given in Figure 1A. Expressed bacteriocins are exported from the host strain using a secretion tag and can then target pathogenic strains in the extracellular environment. Four secretion tags were tested to determine if the level of antimicrobial activity differed between each tag. The bacteriocin coding sequence linked to the secretion tag is under the control of a constitutive promoter.

**Figure 1:**
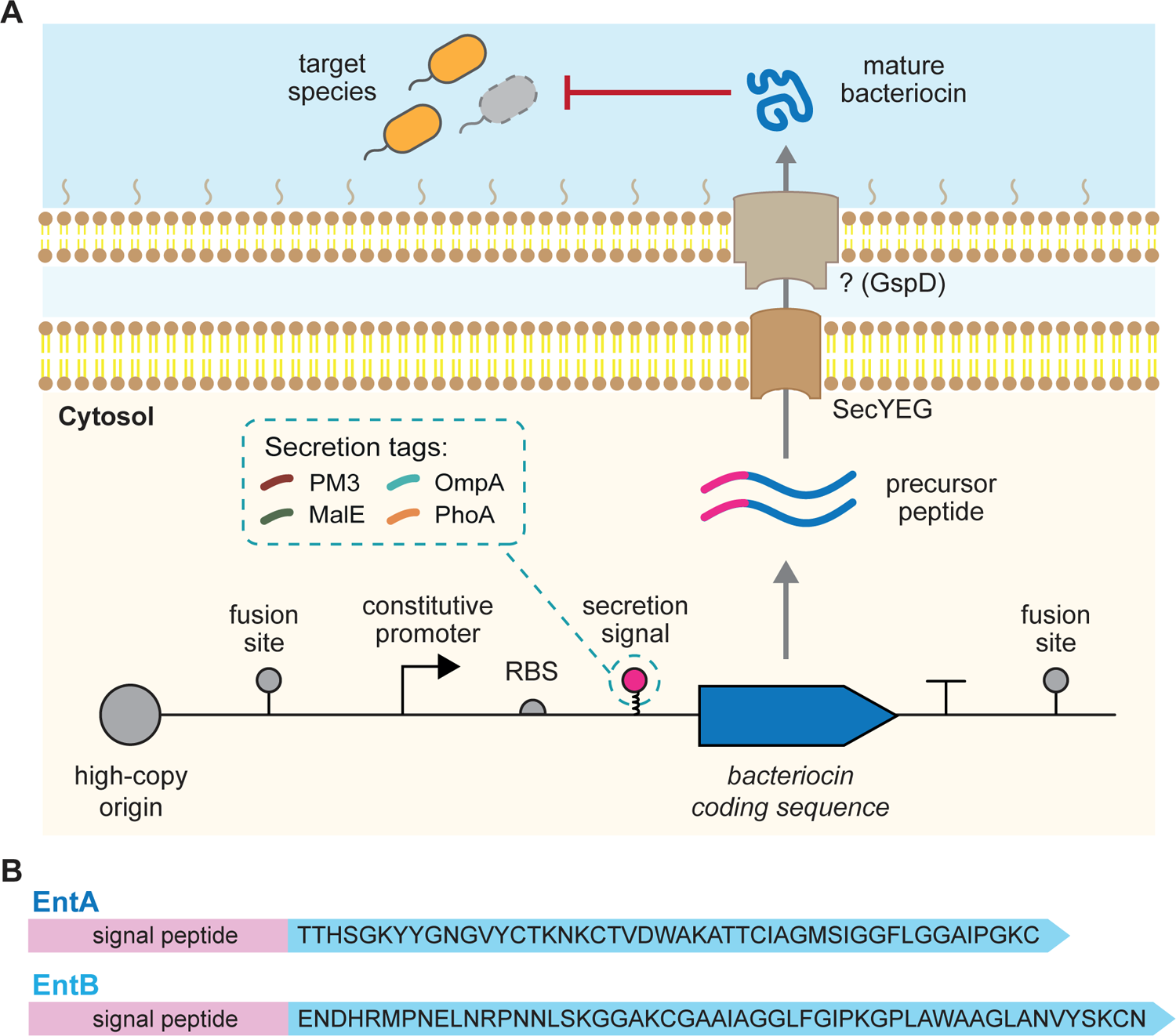
Overview of the bacteriocin expression platform. (A) A high-copy plasmid codes for constitutive expression of a specific bacteriocin fused with one of four different secretion signals. The expressed bacteriocins are secreted from the host cells and can target susceptible species in the extracellular environment. The secretion singals used in this study have previously been reported to exit the cell via the SecYEG inner-membrane and the GspD outer-membrane pore. (B) The amino acid sequences of the EntA and EntB bacteriocins used in this study.

Enterocin A (EntA) and Enterocin B (EntB) were chosen as candidate bacteriocins to show the efficacy of our AMP system. The peptide sequences for both are given in Figure 1B. The antimicrobial activity of these bacteriocins against *E. faecalis* was confirmed via chemically synthesised peptides (SI Figure S1). We also confirmed that EntA and EntB did not show antimicrobial activity against *E. coli* NEB^®^express cells, when exposed to 4*µ*g of synthetic bacteriocin (SI Figure S1C). Growth curves of *E. faecalis* cultures grown in the presence of EntA, EntB or a mixture of both (EntAB) showed that EntAB was able to delay the development of resistance at higher concentrations (SI Figure S1D).

### 2.2 Screening of bacteriocin secretion constructs

We tested the efficacy of our platform against *E. faecalis* in both solid and liquid culture assays. The sequences of the four secretion signals are given in Figure 2A. The predicted cleavage site of these secretion tags was confirmed using the online SignalP 6.0 tool (SI Figure S2)(*36*). An inhibition assay was used to confirm the antimicrobial activity of our engineered strains against *E. faecalis*. All of the engineered strains were found to produce a zone of inhibition. The largest inhibition zone was found with the OmpA-EntA strain and the smallest with the PhoA-EntB strain. No inhibition zone was observed for the control construct which did not contain a bacteriocin expression plasmid (Figure 2C and 2D).

**Figure 2:**
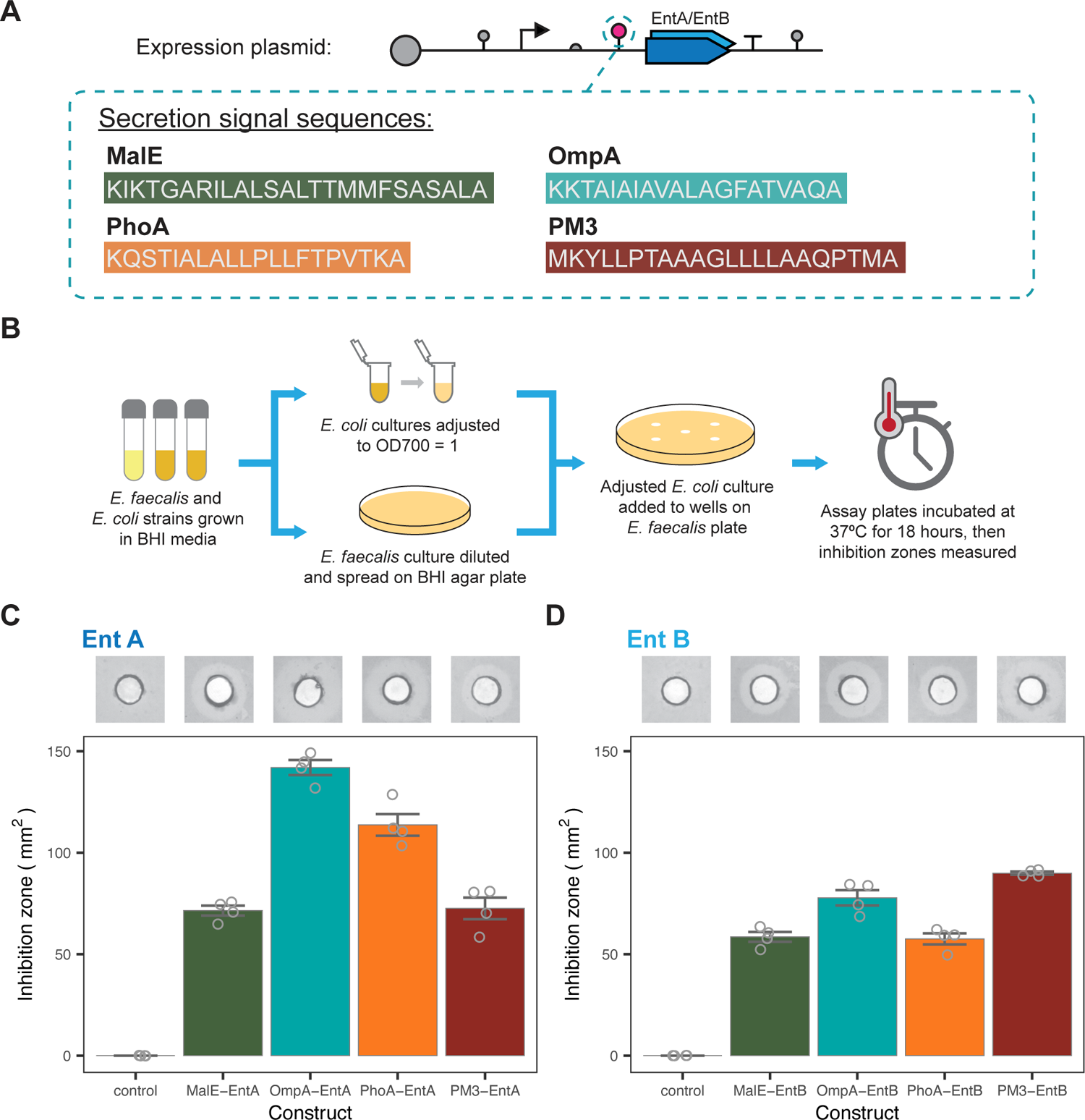
Inhibition zone characterisation of the individual bacteriocin expressing strains. (A) The amino acid sequences of the four secretion signals screened in this study. (B) The experimental protocol used for solid culture inhibition assays. The measured inhibition zones for EntA (C) and EntB (D) secreting strains with all four secretion tags. The control indicates the host strain with no bacteriocin expression plasmid. Insets show representative images of the inhibition zones (central area indicates loading well). Bars indicate mean values ± SE of four individual repeats.

Competition assays between our engineered *E. coli* and *E. faecalis* were performed in liquid culture (Figure 3A). Co-culture dynamics were extracted from bulk measurements using a fluorescent label in the *E. coli* strain (SI Figure S3)(*37*). The competitive exclusion of *E. coli* by *E. faecalis*, seen with the control, is counteracted, with varying degrees of efficacy, by the secretion of bacteriocins (Figure 3B). In order to rank the performance of the strains the proportion of *E. faecalis* at 10 hours was chosen as a summary statistic (Figure 3C). All bacteriocin secreting strains suppressed the growth of *E. faecalis* in comparison with the control strain. In addition, all EntA secreting strains performed better than their EntB counterparts, agreeing with the activity of the synthesised bacteriocins (SI Figure S1) and previous studies that suggest EntA has greater antimicrobial activity(*24*). A comparison of the inhibition zones and co-culture ratio showed that the size of the inhibition zone for each strain was not a good indicator of the killing levels seen in co-culture (Figure 3D).

**Figure 3:**
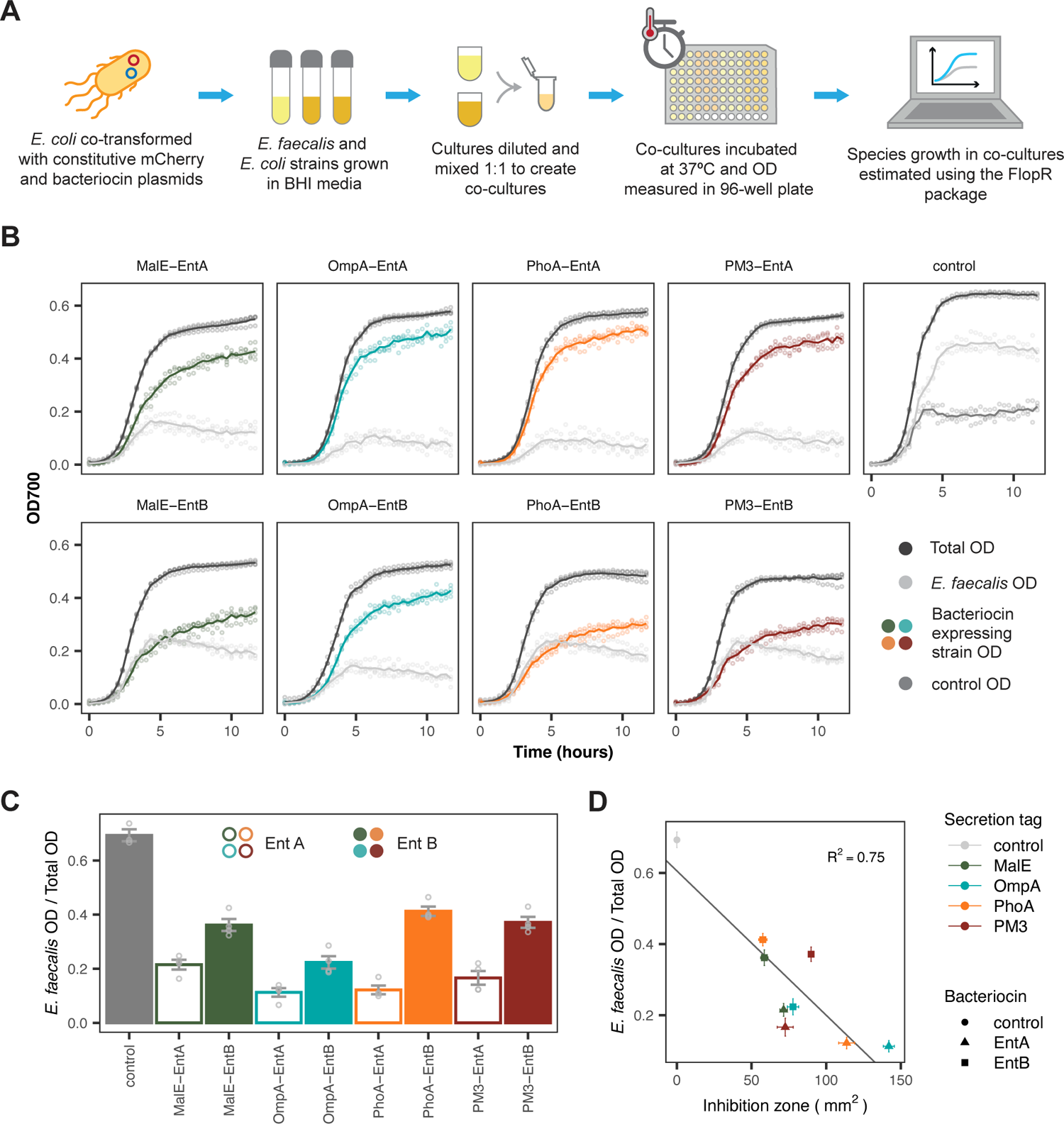
Liquid co-culture characterisation of the individual bacteriocin expressing strains. (A) The experimental protocol used for co-culture growth assays. (B) Growth curves of the competitor strains in co-culture. The labels indicate the bacteriocin construct produced by the engineered competitor strain. Lines indicate mean of four replicates and points show individual repeats. (C) The estimated ratio of *E. faecalis* OD over total OD at 10 hours, when grown in co-culture with the indicated strain. Bars indicate mean values ± SE of four individual repeats. (D) Comparison of the solid culture inhibition zone vs the 10 hour ratio for each of the co-culture conditions. Solid line shows linear regression fit, with the R^2^ value labelled. Points give mean value ± SE of four individual repeats.

We further investigated the affects of initial culture density and *E. faecalis*:*E. coli* (Ef:Ec) seeding ratios using the PM3-EntA secreting strain. Decreasing the initial culture density and increasing the Ef:Ec seeding ratio resulted in higher levels of *E. faecalis* growth (SI Figure S5C). We hypothesise this may be due to the need to reach a threshold level of bacteriocin that can overcome competitive exclusion, which favours the faster growing *E. faecalis* strain. Also, the *E. faecalis* proportion remained relatively stable from six to ten hours (SI Figure S5A and S5B); suggesting that for the conditions tested, final co-culture composition is determined before the co-culture reaches the stationary phase of growth.

### 2.3 Impact of dual-bacteriocin secretion

Next, we set out to explore the effect of secreting multiple bacteriocins simultaneously, as we hypothesised that this will reduce the rate of resistance occurring. Using our plasmid system we constructed four EntAB co-expressing strains that used the same secretion tags for each bacteriocin (Figure 4A). The co-culture assays, once again, demonstrate that the EntAB expressing strains are able to overcome competitive exclusion and suppress *E. faecalis* growth (Figure 4C). The proportion of *E. faecalis* in the co-cultures at 10 hours is less in the MalE-EntAB and PM3-EntAB dual-bacteriocin assays compared to the single bacteriocin assays (Figure 4D), demonstrating that expression of both bacteriocins improves killing for these strains.

**Figure 4:**
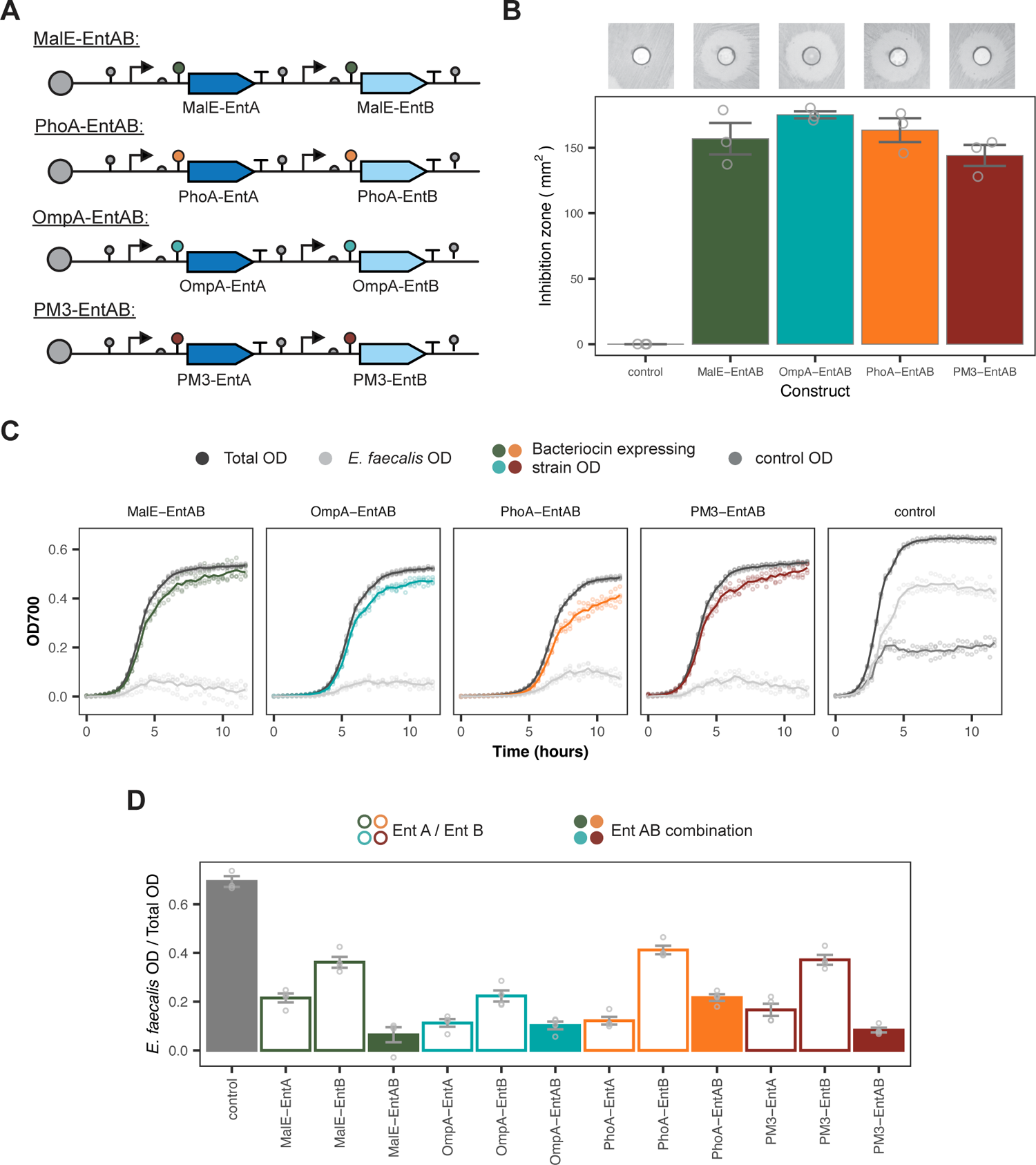
Full characterisation of the dual-bacteriocin expressing strains. (A) Plasmid layout of the co-expressing strains. (B) Inhibition zones of the dual-secreting strains. Insets show representative images of the inhibition zones. Bars indicate mean values ± SE of three individual repeats. (C) Growth curves of strains in co-culture. Labels give bacteriocin construct produced by competitor strain. Lines indicate mean of four replicates and points show individual repeats. (D) Estimated ratio of *E. faecalis* OD over total OD at 10 hours for the dual-bacteriocin strains compared to the ratios given in Figure 3C, when grown in co-culture. Bars give mean values ± SE of four individual repeats.

To check that bacteriocin production did not significantly increase the metabolic burden on the host strain, the growth curves of each strain in monoculture (SI Figure S5A) were fit with a Gompertz model (SI Figure S5B). As expected, the control showed the fastest growth rate of the engineered *E. coli* strains. The control contains only a single fluorescent plasmid and therefore does not have to divert cellular resources for bacteriocin production. Of the bacteriocin producing strains, OmpA-EntAB and PhoA-EntAB were found to have the slowest growth rates and longest lag times (SI Figures S5C and S5E, respectively).

To show the wider application of our system we screened the PM3-EntA expression plasmid in three alternative host strains (*E. coli* BW25113, NEB^®^5-α and Nissle 1917). *E. faecalis* killing was seen at similar levels for all host strains (Figure 5A). In addition, our system showed antimicrobial activity against a vancomycin-resistant isolate of *Enterococcus faecium* (Figures 5B and 5C)(*33*). As with other antimicrobials, bacteria can develop resistance to bacteriocins over time. We used a 12 hour passage experiment to test the development of *E. faecalis* resistance to our PM3-engineered strains. For all PM3-strains, we found that *E. faecalis* was able to out compete the engineered strains after 48 hours in co-culture (SI Figure S6).

**Figure 5:**
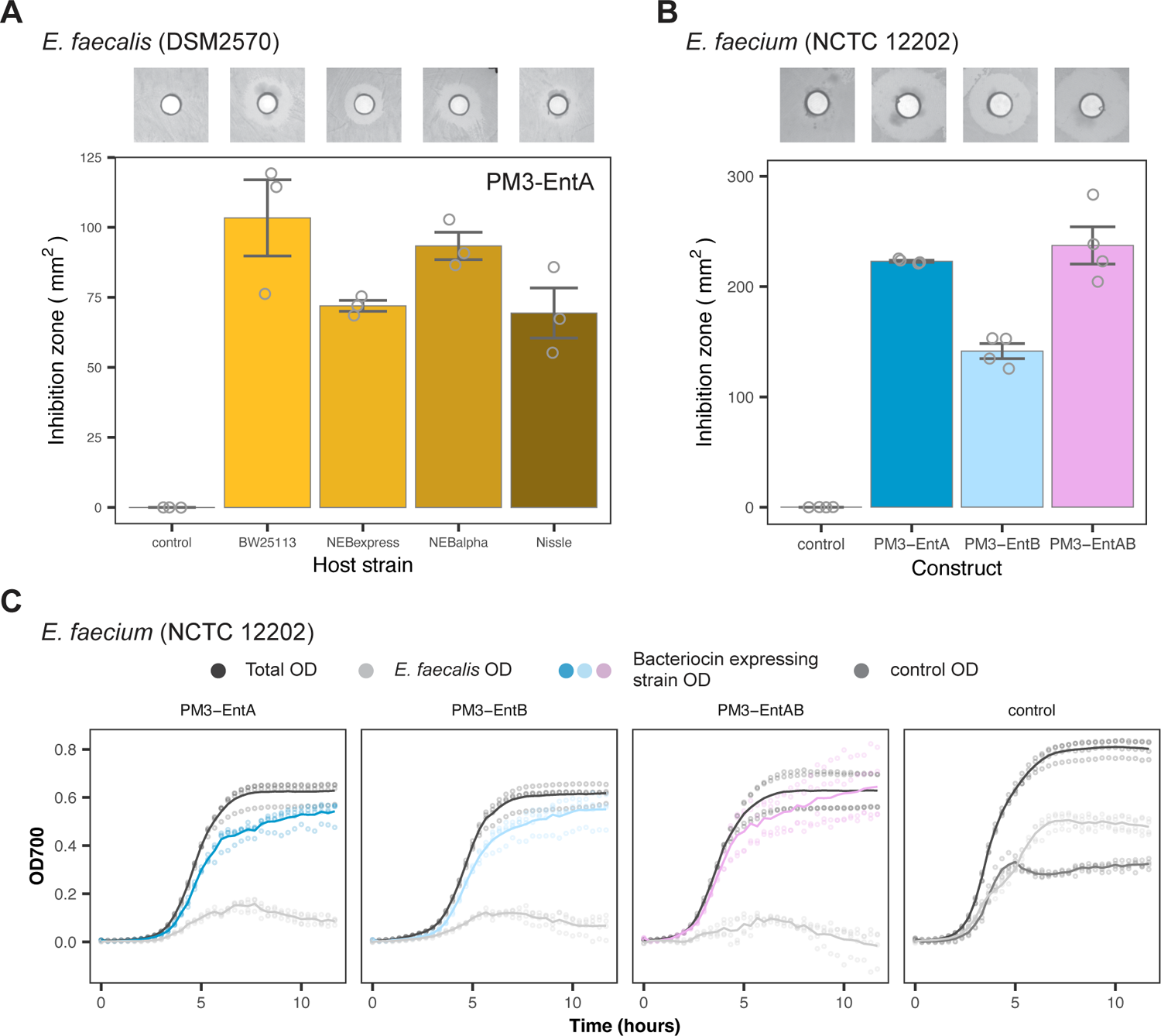
Wider application of the expression system, in alternative hosts and targeting vancomycin-resistant *E. faecium*. (A) Inhibition zone assay of the PM3-EntA construct in multiple *E. coli* host strains. Insets show representative images of the inhibition zones observed (n = 3, bars indicate mean ± SE). (B) Inhibition zone assays for the given constructs against a Vancomyicn-resistant *E. faecium* isolate (n = 4, bars indicate mean ± SE). (C) The growth curves of *E. faecium* grown in co-culture with the labelled competitor strain (n = 4, solid lines give mean values and points individual replicates).

### 2.4 Modelling co-culture dynamics

We used mechanistic modelling and Bayesian statistics in order to gain further insights into the dynamics of the bacteriocin interaction. We jointly fitted the data from the PM3 system including monocultures of *E. faecalis* and *E. coli* single and dual bacteriocin production strains in co-culture with *E. faecalis* (extended SI Methods). Five different models were developed in total. Three of the models were based on a linear pairwise bacteriocin interaction and two on a saturated pairwise interaction (*38*). We used Bayesian model selection to select the best fitting model (SI Figure S7A). The saturated model for bacteriocin interaction fitted the data best with parameters that allows for reduced growth rate and carrying capacity in the bacteriocin producing strains (Figures 6A and 6B).

**Figure 6:**
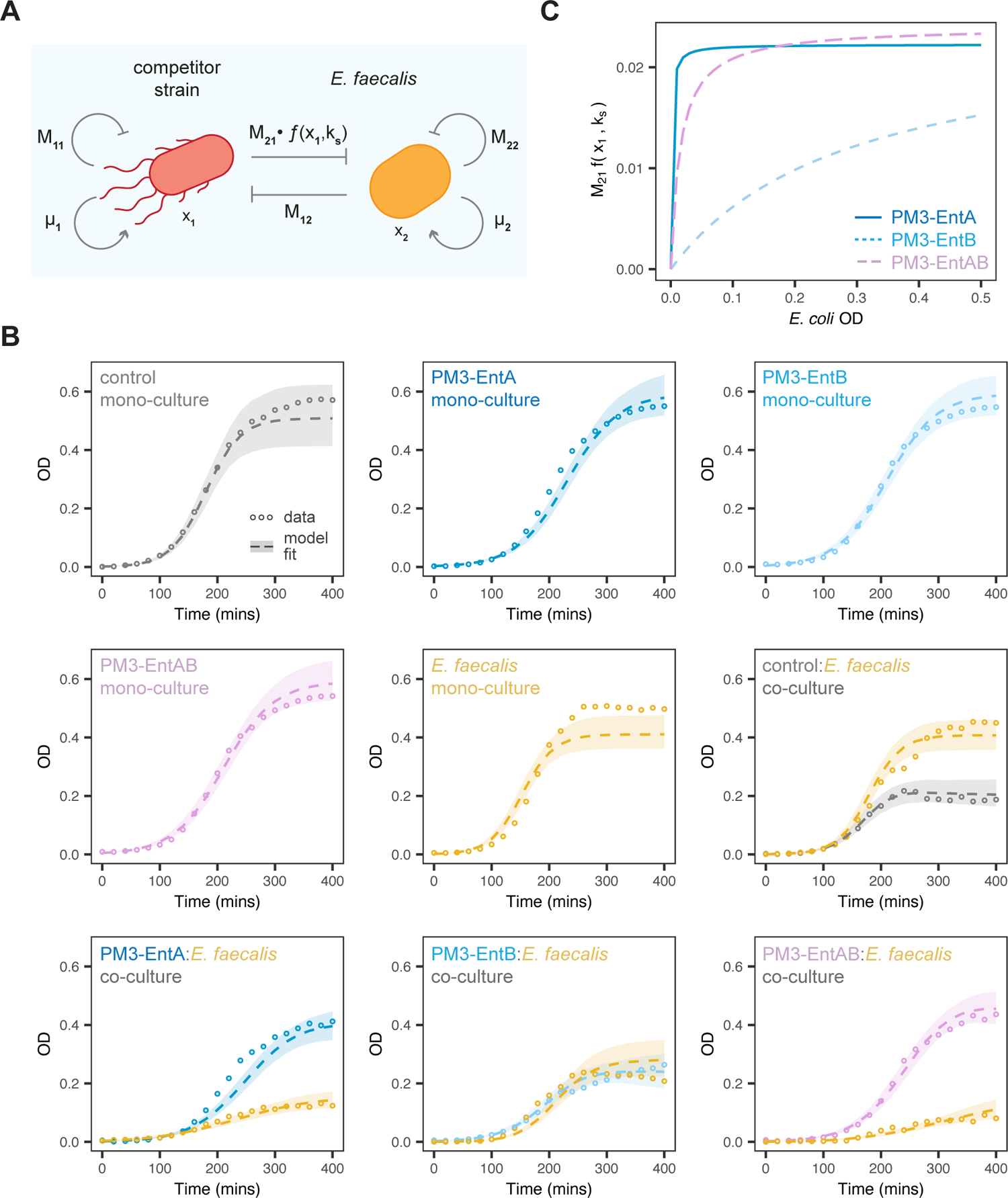
Lokta-Volterra model fitting of co-culture growth dynamics. (A) Schematic of the major model parameters, *x_i_* refers to species, *µ_i_* to species growth rate and *M_ij_*to species interactions. (B) Fitted growth curves for the labelled mono- and co-cultures (circles give mean experimental value, dashed lines give the median simulated timecourse and shaded regions the 95% credible region). (C) The fitted interaction term (*M*_21_·*f* (*x*_1_*, k_s_*)) against the estimated OD for the given strains, using the median of the posterior distribution.

The posterior parameters from the best model (SI Figure S7B) show that *E. faecalis* exerts a competitive effect on *E. coli* so they do have overlapping niches in this medium indicated by a negative *M*_12_. The maximum killing, *M*_21_, has a median value of −0.0223, −0.0242, and −0.0240, for EntA, EntB, EntAB respectively, although the posteriors overlap somewhat due to low information in this experimental setup. The main difference (especially between EntA and EntB) is the *K_s_* value, which is the point of half maximum in the interaction curve. The posterior median curves for the inferred saturation function show bacteriocin killing quickly reaching a maximum in the case of EntA (Figure 6C). This is consistent with SI Figure S1B where there is no noticeable killing until a concentration of 2*µg* of EntB. The dual system, EntAB, shows a behaviour inbetween the individual bacteriocins, with a slightly slower turn on than EntA but a slightly higher maximum (Figure 5C). It was also observed that, because the saturation curves for EntA/EntAB are sharp, the effect on *E. faecalis* is approximately independent of the concentration of *E. coli*. This gives rise to a lower effective *E. faecalis* growth rate, which accounts for the qualitative difference between the EntA, EntAB and the EntB co-culture dynamics.

### 2.5 eLBP performance in a three-strain community

We further tested the activity of our bacteriocin-producing eLBPs in a three-strain community. As the third strain we used a GFP-fluorescent *E. coli* NEB^®^5-α strain, referred to here as the “bystander” species. For the control:*E. faecalis*:bystander community, *E. faecalis* was seen to outgrow the other strains in co-culture (Figure 7B). However, in the presence of the PM3-EntA eLBP, *E. faecalis* growth was suppressed and bystander growth dominated the co-culture. In two-strain co-culture the *E. faecalis* strain was able to out-compete the bystander (Figure 7C).

**Figure 7:**
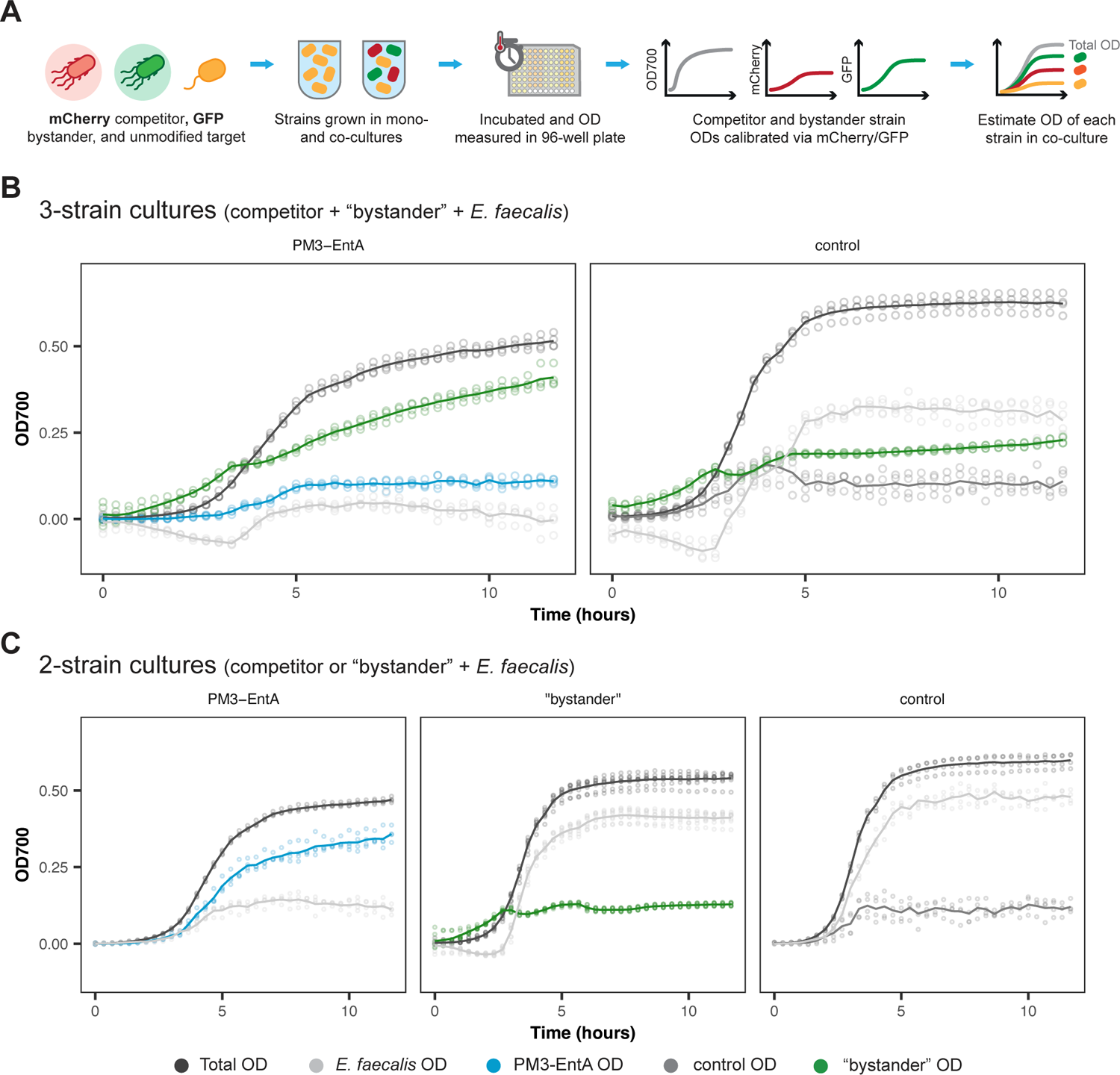
Three-strain community culture dynamics. (A) Overview of the experimental process for measuring growth in the three-strain communities. GFP and mCherry fluorescent signals are used to estimate the OD of the “bystander” and competitor strains, respectively. (B) Growth curves of the PM3-EntA competitor, bystander and *E. faecalis* (left panel) and the control competitor, bystander and *E. faecalis* (right panel) strains in co-culture. (C) Growth curves of the PM3-EntA, bystander or control strain in co-culture with the *E. faecalis* target strain in two-strain co=culture. Lines indicate mean of four replicates and points show individual repeats.

## 3 Discussion

The primary goal of this study was to create a novel AMP production platform that could be used to express bacteriocins from a commensal host. We built on previous works by developing a modular platform based on the CIDAR MoClo assembly standard(*34*). Unlike previous *ad hoc* systems, our platform is based on a previously established cloning standard and therefore, can make use of a wide range of pre-existing parts. These include inducible promoters and biosensors, which allow for the tailoring of bacteriocin expression towards specific scenarios. Our system allows for the rapid construction of strains that are able to produce bacteriocins from separate operons. Not only does this help remove the variable of operon layout, it also allows for modification of expression levels based on interchangeable promoters and RBSs. We also screened four secretion signals: MalE, OmpA, PhoA and PM3. These tags were chosen as they are compatible with *E. coli*, have been shown to express proteins in heterologous systems, and use the general secretion (Sec) pathway(*39*). The Sec pathway offers high export capacity and a lack of specificity that is desirable for a platform that can be used to secrete a range of different bacteriocins(*17*). It is worth noting that the four secretion tags explored here are generally reported as targeting protein secretion to the periplasm. A previous study suggested that proteins expressed with the PelB secretion signal are subsequently exported from the periplasm via the GspD secretion pore(*40*). To confirm whether this was the case for our PM3-EntA secreting strain, a modified version of the PM3-EntA plasmid was developed including constitutive expression of the *gspD* gene (termed PM3-EntA-GspD). These constructs were then screened in both a *gspD* knockout and parent strain chassis. Even for strains with no GspD expression, killing zones were seen (SI Figure S8), suggesting that the GspD pore is not solely responsible for bacteriocin secretion from these strains. This will need to be explored further, to identify which secretion pores are responsible for the killing seen from our engineered strains.

Typically the performance of AMP producing strains is quantified through supernatant growth assays that measure the growth of the target strain when exposed to either conditioned supernatant or purified AMP(*23*, *24*, *41*). However, these assays do not reflect the mode of delivery in which these engineered strains are designed to operate(*22*). One of the proposed benefits of using engineered bacteria is that the AMP can be expressed directly at the desired site within the body. Within this scenario the AMP secreting and target strains will likely be growing in close proximity. To more closely mimic these operating conditions we used co-culture assays to assess the performance of our bacteriocin secreting strains. We used our existing tool, FlopR, for the rapid determination of antimicrobial activity of species grown in co-culture. As the FlopR package only requires one species in a two-species co-culture to be fluorescent, it is feasible to rapidly screen antimicrobial activity against multiple strains without the need for any modification of the target species(*37*).

However, under typical *in vivo* conditions other strains of the host microbiota will be present in the environment, alongside the engineered and target strains. Therefore, to further demonstrate the utility of our FlopR pipeline we characterised the PM3-EntA eLBP and control strains in a three-species community. In theory, FlopR analysis can be extended to an *n* number of species co-culture; provided that *n* − 1 distinct fluorescent signals can be measured. For the three-strain communities GFP and mCherry signals were used to estimate the growth of the bystander and competitor strains, respectively. To achieve this we engineered a GFP expressing bystander and confirmed that other species in the co-culture did not produce a GFP signal (SI Figure S9). In two-strain co-culture *E. faecalis* was able to outgrow the bystander strain (Figure 7C). This was also the case in the control:bystander:*E. faecalis* co-culture. In contrast, in the PM3-EntA:bystander:*E. faecalis* co-culture, *E. faecalis* growth was suppressed and the bystander strain was able to dominate the measured OD (Figure 7B). This suggests that the PM3-EntA eLBP strain is able to suppress growth of the target strain, creating a niche that the bystander strain is able to exploit.

In addition, anaerobic inhibition zone assays were performed to confirm antimicrobial activity of the PM3-bacteriocin strains in anaerobic environments (such as those found within the digestive tract). All three of the bacteriocin expressing strains tested produced inhibition zones against *E. faecalis* (SI Figure S10).

Previous studies have shown that administering bacteriocins in combination can result in improved antimicrobial activity(*24*). In addition, it has been reported that when co-expressed EntA and EntB can form a synergistic heterodimer(*42*). The MalE-EntAB and PM3-EntAB strains suppressed *E. faecalis* growth more than the individual EntA/EntB counterparts (Figure 4). However, OmpA-EntAB and PhoA-EntAB did not improve on the activity shown in the single bacteriocin OmpA-EntA/PhoA-EntA strains in co-culture. This may have been due to the increased metabolic burden placed on the hosts of these constructs, which is indicated by the decreased growth rate and increased lag time of these strains (SI Figure S5)(*43*).

Finally, we used mathematical modelling and Bayesian statistics to gain insight into the dynamics using the co-culture data. This is important because in the case of eLBPs there will be a trade-off between production of AMP and metabolic burden, with subsequent lower growth rate and reduced fitness. We tested a number of pairwise interaction models and compared how well they fit using Bayesian model selection (Figure 6, Figure S7). A model containing a saturated Lotka-Volterra term for bacteriocin interaction was the best model. Although this a common pairwise model it can also be derived from the full mechanistic model assuming that either a) the bacteriocin producing strain grows faster than the target or b) that the bacteriocin is reusable (*38*). Since *E. faecalis* grows faster than *E. coli* we can assume the model implies that the bacteriocin is reused after killing. We gained a number of other insights into the dynamics, including the form of the saturation functions for the different bacteriocin systems, that EntB has a much higher *K_s_* value in the saturation function than EntA, and quantification of the metabolic burden of bacteriocin expression. We anticipate that as more complex eLBPs are produced over the coming years it will be increasingly important to characterise these dynamics using mathematical models.

To date we have only screened EntAB co-expressing strains that use the same secretion signal. However, using multiplex reactions it is feasible to create strains that use a mixture of secretion signals. It is possible that other combinations may be able to alleviate negative growth impacts on the co-expressing strains while also increasing antimicrobial activity. We advocate using co-cultures and mathematical modelling to assess eLBP performance. However, even in co-culture, the conditions found within the human gut will be extremely different to those encountered *in vitro*. Therefore, *in vivo* testing or improved *in vitro* methods will be required to more accurately predict final strain performance(*44*). We also measured co-culture dynamics only indirectly using FlopR. To confirm the estimated reduction in *E. faecalis* growth shown in the FlopR assays was correct, an assay was developed in which the eight hour timepoint of co-culture growth was sampled and used to estimate colony counts of *E. faecalis* growth when grown with the PM3-bacteriocin secreting strains. This assay confirmed that the EntA and EntAB constructs reduced *E. faecalis* growth in comparison to the control strain (see SI Figure S11). As was seen in the FlopR assays, the EntB construct had a much smaller impact on *E. faecalis* growth. Although we used pairwise mathematical models to interrogate the co-culture dynamics, mechanistic models where bacteriocin concentration is explicitly modelled would provide separate information on production and killing. Although measuring bacteriocin concentration over time in a high-throughput manner is probably infeasible, more information could be leveraged by combining co-cultures with synthetic bacteriocin spike-in experiments. Combined with Bayesian hierarchical modelling, this could allow separation of the production and killing rates.

In conclusion, we have developed a modular bacteriocin secreting system that can be used to suppress the growth of *E. faecalis* and VRE. Although we focussed on *Enterococcus* species, the modular nature of our system can make use of expanding libraries of bacteriocins (*45*) to design novel antimicrobials specifically targeting other species. Additionally, our experimental approach can be applied to the rapid characterisation of these antimicrobial-secreting eLBPs. In future, we will also leverage multiplexed assembly reactions afforded by our CIDAR MoClo based system(*34*), to screen for bacteriocin constructs with the greatest antimicrobial activity. eLBPs like those developed here could be an important tool against the global increase in antimicrobial resistant pathogens.

## 4 Methods

### Strains and plasmids

Construct characterisation was performed in Brain Heart Infusion (BHI) media (Sigma Aldrich). All secretion signal-bacteriocin sequences were purchased as gBlock fragments (IDT), with any internal Bsa-I and Bbs-I restriction sites removed. All full bacteriocin secretion plasmids were constructed using DNA parts from the standard CIDAR MoClo kit(*34*). All bacteriocin expressing plasmid transcription units within this work were constructed with the J23106 constitutive promoter, BCD12 RBS and B0015 terminator. Cloning was performed in commercial NEB^®^5-α competent *E. coli* (New England Biolabs) and plasmid sequences confirmed with Sanger sequencing. Correct plasmids were then transformed into the desired competitor strains and stored as glycerol stocks at −70^◦^C until needed. Unless otherwise stated all characterisation experiments were performed with commercial *E. coli* NEB^®^express host cells. The target, *E. faecalis* DSM2570, was purchased from DSMZ-German Collection of Microorganisms and Cell Cultures GmbH. A full list of the strains and plasmids used in this study is given in SI Table 1.

### Solid culture characterisation

Bacteriocin secreting strains and *E. faecalis* were inoculated from glycerol stocks and grown in BHI media for ∼18 hours at 37^◦^C with shaking. Two *µ*Ls of the *E. faecalis* culture was then diluted in 150 *µ*L of fresh BHI media and spread on a 30 mL 1.5% BHI agar plate and allowed to dry. Cultures of the bacteriocin secreting strains were then adjusted to OD_700_ 1.0 in sterile BHI media. Loading wells were stamped in the dried BHI agar plates using a sterile glass Pasteur pipette. Adjusted culture (50 *µ*L) was added to the loading wells and the plates incubated for ∼18 hours at 37^◦^C. Assay plates were then imaged using a LoopBio Imager and inhibition zones measured manually in FIJI software(*46*). The inhibition zone was defined as the area that showed no *E. faecalis* growth minus the area of the loading well.

### Liquid co-culture characterisation

Bacteriocin secreting strains were co-transformed with a constitutive mCherry2 fluorescent plasmid (built with the J23101 promoter and p15a origin). All co-transformed strains and *E. faecalis* were inoculated from glycerol stocks and grown in BHI media for ∼18 hours at 37^◦^C with shaking. Cultures were then adjusted to OD_700_ 1.0 and diluted 100-fold in fresh BHI media. For monocultures, 120 *µ*L of each diluted culture was added directly to the well of a 96-well plate. For co-cultures, the desired ratio of each strain was added (up to a total of 120 *µ*L) to each well of a 96-well plate. Plates were then incubated for 24 hours at 37^◦^C with shaking (300 rpm, 2 mm orbital), in a Tecan Spark plate reader (Tecan, USA). Measurements for OD_700_ and mCherry fluorescence (excitation: 531/20 nm, emission: 620/20 nm, gain: 120) were collected every 20 minutes.

The growth of each individual strain in the co-culture was then estimated using the FlopR package. A detailed description of this process is given by Fedorec *et al* (2020)(*37*). In brief, a calibration curve of the fluorescence in each mono-culture was used to estimate the fraction of fluorescent cells in each corresponding co-culture, based on the ratio of expected vs measured fluorescence for a given OD measurement. This fraction was then used to estimate the change in species abundance across time in each co-culture. This process is summarised in SI figure S3.

## Data analysis and visualisation

All data analysis and visualisation was performed in Rstudio (version R4.1.2). Further details of the materials and methods can be found in the Supporting Information.

## Data availability

The data and code to reproduce the results in Figures 2 to 4 can be found in a Zenodo repository: doi 10.5281/zenodo.8427304.

## Acknowledgement

The authors thank Michael J Bland and Philippe Gabant (Syngulon SA, Belgium) for kindly providing chemically synthesised EntA and EntB bacteriocin samples.

## Supporting Information Available

Supplementary Figures S1-S11 and extended materials and methods.

