## Supplementary information for "An antimicrobial peptide expression platform for targeting pathogenic bacterial species"

**2 Extended results** **7**

**References** **19**

### 1 Extended materials & methods

#### 1.1 Strains & plasmids

All strains and plasmids used in this study are given in Table 1.

#### 1.2 Synthetic bacteriocin assays

Assay plates were prepared with 30 ml of sterile BHI 1.5% agar in 90 mm petri dishes. Plates were allowed to set before 2 $\mu$l of overnight culture for the target strain was diluted into 150 $\mu$l of fresh BHI and spread on the surface of the assay plate. Once dry, loading wells were cut into the agar using a sterile glass Pasteur pipette. 50 $\mu$l of desired synthetic bacteriocin concentrations were then added to the loading wells and the plates incubated for 18 hours at 37°C. Images were then collected and processed as described in the main text.

Synthetic bacteriocin timecourses were performed on *E. faecalis* mono-cultures. Overnight *E. faecalis* cultures were adjusted to OD700 = 1 and then diluted 1:100 in fresh BHI media. The diluted cultures were then supplemented with the desired bacteriocin concentrations and 120$\mu$L added to each well of a 96 well plate. The growthcurves were then recorded by measuring OD700 absorbance every 20 minutes for a total of 48 hours.

#### 1.3 GspD knockout assays

The ΔgspD knockout strain was purchased from the Keio knockout collection[1] and transformed with the desired bacteriocin expression plasmid. The inhibition zone assays was performed as detailed in the standard solid culture characterisation protocol given in the main text.

#### 1.4 Three-strain co-culture assays

Liquid culture assays were performed as described in the main text, with the GFP fluorescent bystander strain added to all three-strain cultures (up to a total of 120$\mu$L). Measurements of GFP (excitation: 488/20 nm, emission: 530/20 nm, gain: 70) fluorescence were collected alongside $OD_{700}$ and mCherry readings.

#### 1.5 Anaerobic inhibition zone assays

Anaerobic inhibition zones assays were performed as described in the main text, with two modifications. Both the 18 hour incubation of inoculated strains and the final 18 hour incubation of the assay plate were performed in an anaerobic environment. Anaerobic conditions were maintained with a miniMACS anaerobic workstation (Don Whitley Scientific, UK).

Table 1: Strains and plasmids used within this study (RBS = ribosome binding site, CDS = coding DNA sequence).

| **Strain** | **Description** | **Source** |
|---|---|---|
| *E. coli* NEB®5-$\alpha$ | Commercial cell line used for cloning | New England Biolabs |
| *E. coli* NEB®Express | Commercial cell line used for protein expression | New England Biolabs |
| *E. coli* Nissle 1917 | Commensal strain of *E. coli* | Prof. Ian Henderson (Uni. of Birmingham, UK) |
| *E. coli* BW25113 | Parent strain of the Keio knockout collection | Keio Collection |
| *E. coli* JW5707 | BW25113 Δ*gspD* knockout strain | Keio Collection |
| *E. faecalis* DSM25700 | *E. faecalis* strain | DSMZ collection |
| *E. faecium* NCTC12202 | Vancomycin-resistant isolate of *E. faecium* | Dr Julie MacDonald (Imperial College, UK) |
| **Plasmid** | **Description** | **Source** |
| pMalE-EntA | J23106 promoter, BCD12 RBS, MalE-EntA CDS, B0015 terminator, DVK_AF vector, Kan$^R$ | This study |
| pMalE-EntB | J23106 promoter, BCD12 RBS, MalE-EntB CDS, B0015 terminator, DVK_FG vector, Kan$^R$ | This study |
| pMalE-EntAB | pMalE-EntA, pMalE-EntB, DVA_AG vector, Amp$^R$ | This study |
| pOmpA-EntA | J23106 promoter, BCD12 RBS, OmpA-EntA CDS, B0015 terminator, DVK_AF vector, Kan$^R$ | This study |
| pOmpA-EntB | J23106 promoter, BCD12 RBS, OmpA-EntB CDS, B0015 terminator, DVK_FG vector, Kan$^R$ | This study |
| pOmpA-EntAB | pOmpA-EntA, pOmpA-EntB, DVA_AG vector, Amp$^R$ | This study |
| pPhoA-EntA | J23106 promoter, BCD12 RBS, PhoA-EntA CDS, B0015 terminator, DVK_AF vector, Kan$^R$ | This study |
| pPhoA-EntB | J23106 promoter, BCD12 RBS, PhoA-EntB CDS, B0015 terminator, DVK_FG vector, Kan$^R$ | This study |
| pPhoA-EntAB | pPhoA-EntA, pPhoA-EntB, DVA_AG vector, Amp$^R$ | This study |
| pPM3-EntA | J23106 promoter, BCD12 RBS, PM3-EntA CDS, B0015 terminator, DVK_AF vector, Kan$^R$ | This study |
| pPM3-EntB | J23106 promoter, BCD12 RBS, PM3-EntB CDS, B0015 terminator, DVK_FG vector, Kan$^R$ | This study |
| pPM3-EntAB | pPM3-EntA, pPM3-EntB, DVA_AG vector, Amp$^R$ | This study |
| pFlopR-mCherry | p15A origin, J23101 promoter, Elowitz strong RBS, mCherry2, ECK120033736 terminator, Strep$^R$ | This study |
| pFlopR-GFP | J23106 promoter, B0032m RBS, SfGFP CDS, B0015 terminator, DVK_AF vector, Kan$^R$ | This study |
| pPM3-EntA(amp) | J23106 promoter, BCD12 RBS, PM3-EntA CDS, B0015 terminator, DVA_AE vector, Amp$^R$ | This study |
| pPM3-EntA-GspD | J23106 promoter, BCD12 RBS, PM3-EntA CDS, B0015 terminator, J23106 promoter, B0032m RBS, GspD CDS, B0015 terminator, DVA_AF vector, Amp$^R$ | This study |

##### 1.6 Colony-counting assays

Co-cultures of the desired strains were set up following the standard liquid co-culture characterisation protocol given in the main text. After eight hours the timecourse was paused and a 1 $\mu$L sample of each culture was serially diluted in fresh BHI media. 5 $\mu$L of each dilution was plated on an *E. faecalis* selection plate (30 ml of BHI media supplemented with 10 $\mu$g/ml gentamycin). These plates were then incubated for approximately 18 hours at 37°C and the resultant growth used to estimate colony forming units.

##### 1.7 Gompertz growthcurve fitting

A Gompertz growth model was used to fit the mono-culture growth curves shown in Figure S5. The fitting was performed using the nls function in R, using equation (1):

$$y = A \cdot e^{-e^{((\mu \cdot e^{1})/A)\cdot(\lambda - x)+1}} \tag{1}$$

Where $x$ is time, $y$ the measured optical density, $\mu$ the predicted growth rate, $\lambda$ the predicted lag time and $A$ the predicted carrying capacity. These parameters are depicted in Figure S5B.

##### 1.8 Pairwise mathematical models and Bayesian analysis

In the following $x_1$ refers to *E. coli* and $x_2$ refers to *E. faecalis.* Since we don't observe the dynamics of the bacteriocins directly we revert to pairwise models to capture the behaviour.

###### 1.8.1 Baseline model: linear Lotka-Volterra

**Control (no bacteriocin production)**
Here we set $M_{21}$ to zero and have non zero $M_{12}$, motivated by the observation that the levels of *E. faecalis* seem independent of whether the control *E. coli* is present.

$$\begin{aligned} \dot{x_1} &= x_1\mu_{Ec} - x_1M_{11}x_1 - x_1M_{12}x_2 \\ \dot{x_2} &= x_2\mu_{Ef} - x_2M_{22}x_2 \end{aligned}$$

**Bacteriocin production**
We assume the same functional form for EntA and EntB producing *E. coli.* Now we include a non zero (negative) $M_{21}$ to capture bacteriocin action on *E. faecalis.*

$$\begin{aligned} \dot{x_1} &= x_1\mu_{A/B/AB} - x_1M_{11}x_1 - x_1M_{12}x_2 \\ \dot{x_2} &= x_2\mu_{Ef} - x_2M_{22}x_2 - x_2M_{21}^{A/B/AB}x_1 \end{aligned}$$

###### 1.8.2 Saturated Lotka-Volterra with reusable bacteriocin

Here we modify the negative interaction due to bacteriocin action to take a saturated form. Although this is a common pairwise model it can also be derived from the full mechanistic model assuming that either $x_1$ grows faster than $x_2$ or the exchange molecule produce by $x_1$ (the bacteriocin) is reusable [2]. Since *E. faecalis* ($x_2$) grows faster than *E. coli* ($x_2$) we can assume the model captures the second scenario.

**Bacteriocin production**

$$\dot{x_1} = x_1\mu_{A/B/AB} - x_1M_{11}x_1 - x_1M_{12}x_2$$
$$\dot{x_2} = x_2\mu_{Ef} - x_2M_{22}x_2 - \frac{x_2M_{21}^{A/B/AB}x_1}{K_s + x_1}$$

##### 1.8.3 Joint modelling of experiments

We assume that:

- The mutational burden of bacteriocin production is the same for EntA and EntB but different to the control strain. This means we consider three growth rates: $\mu_{Ec}$, $\mu_B$, $\mu_{Ef}$.
- $M_{22}$ is the same in each co-culture experiment. That is, *E. faecalis* self interaction doesn't depend on bacteriocin production.
- The interaction between *E. faecalis* on *E. coli*, $M_{12}$, to be the same across experiments.

We explored how $M_{11}$, self interaction of *E. coli*, changes between control and engineered strains using Bayesian model selection.

##### 1.8.4 Models explored

**M1** : Linear Lotka-Volterra with two self interaction terms: $M_{11}$ and $M_{22}$ for *E. coli* and *E. faecalis* respectively.

**M2** : Linear Lotka-Volterra with five self interaction terms: $M_{22}$ *E. faecalis* and four different $M_{11}$ for the individual *E. coli* strains.

**M3** : Linear Lotka-Volterra with three self interaction terms: $M_{22}$ *E. faecalis* and two different $M_{11}$ for the control and engineered *E. coli* strains (same terms for EntA, EntB and EntAB producing strains).

**M4** : Saturated Lotka-Volterra (reusable bacteriocin) with two self interaction terms: $M_{11}$ and $M_{22}$ for *E. coli* and *E. faecalis* respectively.

**M5** : Saturated Lotka-Volterra (reusable bacteriocin) with three self interaction terms: $M_{22}$ *E. faecalis* and two different $M_{11}$ for the control and engineered *E. coli* strains (same terms for EntA, EntB and EntAB producing strains).

##### 1.8.5 Statistical model

**Likelihood** We assume a LogNormal likelihood, which is equivalent to fitting on the log scale with normally distributed errors:

$$x_i(t) \sim \text{LogNormal}(\log(\hat{x}_i(t)), \sigma)$$

where $x_i(t)$ is the observed data of species $i = 1, 2$ and $\hat{x_i}(t)$ is corresponding output from the ODE model. We simultaneously fit the mono and co culture data, comprising nine datasets: *E. coli* control, *E. coli* EntA, *E. coli* EntB, *E. coli* EntAB, *E. faecalis*, *E. coli* control vs *E. faecalis*, *E. coli* EntA vs *E. faecalis*, *E. coli* EntB vs *E. faecalis*, *E. coli* EntAB vs *E. faecalis*.

**Priors**

$$\begin{aligned}
\mu &\sim \mathrm{U}(0, 0.05) \\
M_{ii} &\sim \mathrm{U}(-0.2, 0) \\
M_{12} &\sim \mathrm{U}(-0.1, 0) \\
M_{21} &\sim \mathrm{U}(-0.2, 0) \\
x(t=0) &\sim \mathrm{LogNormal}(\log(0.01), 1.0) \\
\sigma &\sim \mathrm{LogNormal}(-1, 1) \\
K_s &\sim N(0.2, 0.1)\mathbb{I}(x > 0)
\end{aligned}$$

We used the Rstan interface [3] to the Stan probabilistic programming language [4] to fit the model. All models were fit using 4 chains and convergence verified using the criterion $\hat{R} < 1.1$. Each chain contained 1000 samples, with the first 500 discarded. The final 500 samples of all four chains were combined for the final posterior estimate. All the code for the fitting is included in the Zenodo repository.

#### 2 Extended results

Figure S1 shows the screening of synthetic EntA and EntB activity against *E. faecalis* and *E. coli* NEB®express cells.

Figure S2 gives predictions of the secretion signal cleavage sites.

Figure S3 summarises the FlopR process used for predicting individual strain growth in co-culture.

Figure S4 provides co-culture heatmaps and timecourses, exploring the affects of initial culture density and seeding ratio on final co-culture composition.

Figures S5 show the mono-culture growth curves and Gompertz parameter fits for all strains used in this study.

Figure S6 shows 48 hour timecourses of *E. faecalis* grown in co-culture with the control, PM3-EntA, PM3-EntB and PM3-EntAB secreting strains.

Figure S7 gives a comparison of all the models tested within this study and provides the posterior distributions of all the parameters fitted within the Lokta-Volterra model.

Figure S8 shows the antimicrobial activity of the PM3-EntA bacteriocin constructs (with and without plasmid expression of the *gspD* gene), when expressed from the $\Delta gspD$ knockout (JW5707) and $gspD^{+}$ (BW25113) host strains.

Figure S9 summarises the GFP fluorescence measurements that were used to calculate the bystander strain OD in the 3-strain co-culture assays.

Figure S10 gives the results of inhibition zone assays for the PM3-bacteriocin expressing strains against *E. faecalis*, when grown in anaerobic conditions.

Figure S11 provides estimates of the *E. faecalis* colony counts taken directly from the FlopR co-cultures after 8 hours of growth.

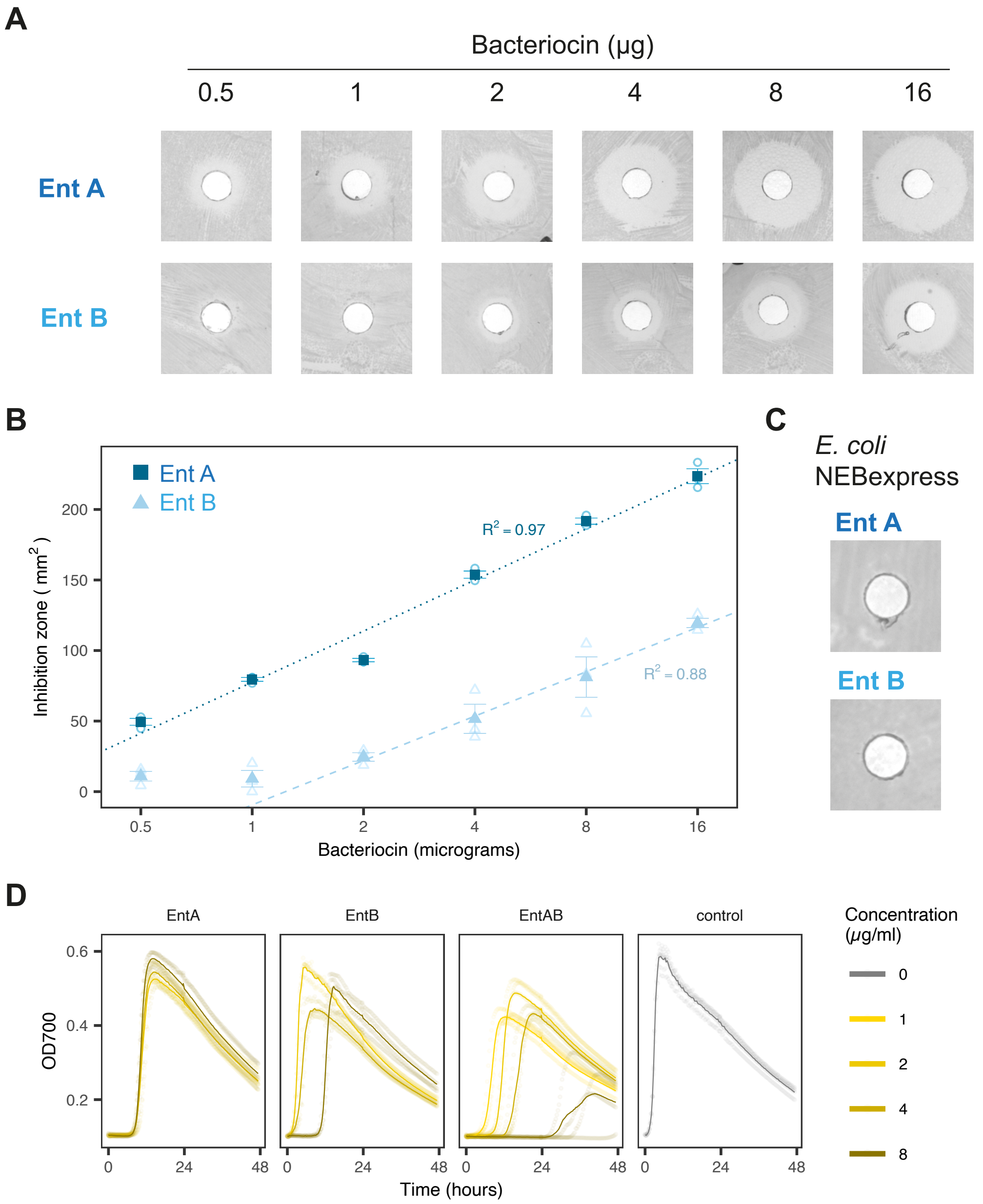

Figure S1: Antimicrobial activity of synthetic Ent A and B, against *E. faecalis*. (**A**) Representative images of the inhibition zones seen when *E. faecalis* was exposed to varying bacteriocin masses. (**B**) Standard curves of Ent A and Ent B inhibition zones for different bacteriocin masses. The dashed lines indicate linear regression fits labelled with $R^2$ values, for Ent B the lowest mass was excluded from the linear regression fit as the zone size appeared to plateau (n = 3, mean ± SE with individual data points). (**C**) Confirmation that no killing was seen for EntA or EntB against an *E. coli* NEB®express lawn. (**D**) Growth curves of *E. faecalis* grown with the given concentrations of synthetic bacteriocin over 48 hours (n = 3, solid lines give mean values).

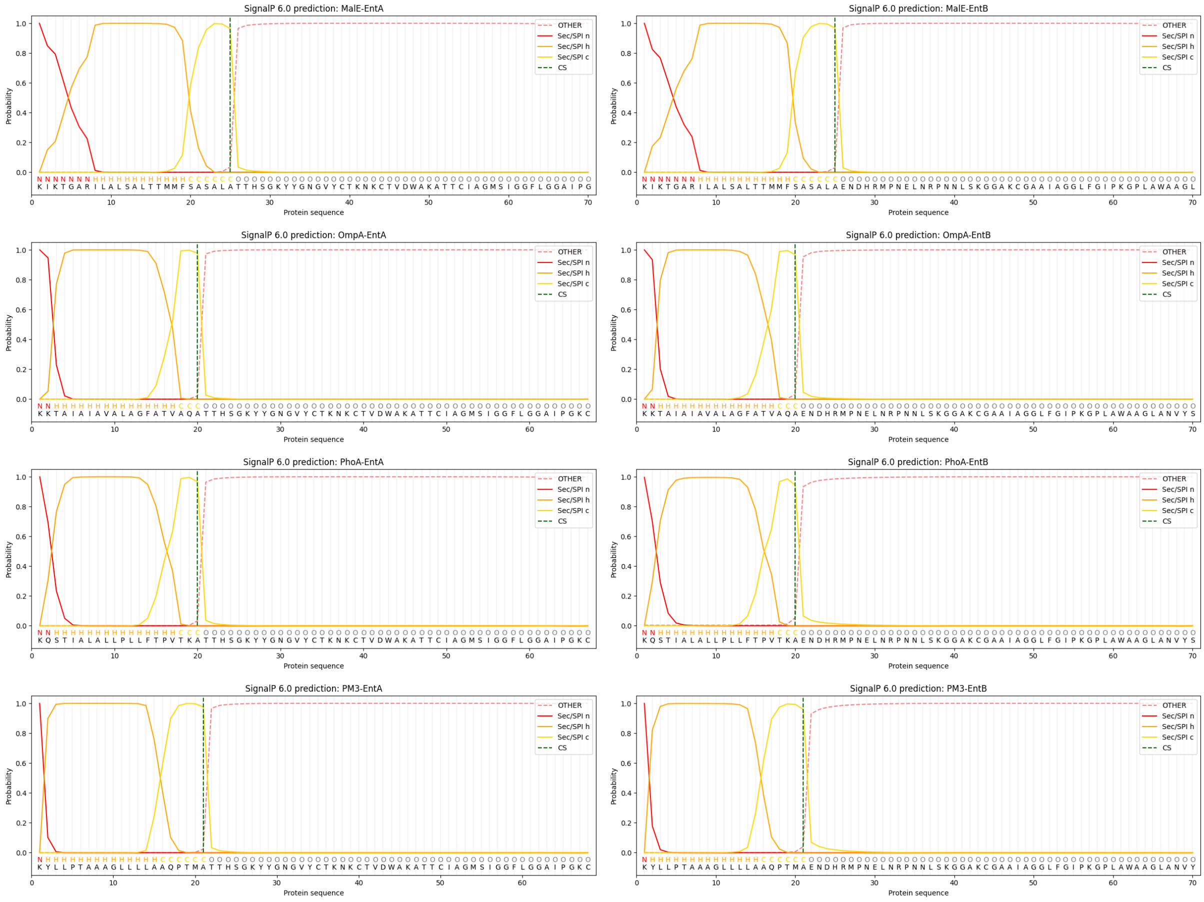

Figure S2: Predictions of the secretion signal locations and cleavage sites, given by the SignalP 6.0 online prediction tool. The predictions are given for all four secretion tags, for EntA (left column) and EntB (right column). SP refers to signal peptide, n denotes the n-region of the peptide, h the h-region and c the c-region. For a full description see Teufel *et al* (2022)[5].

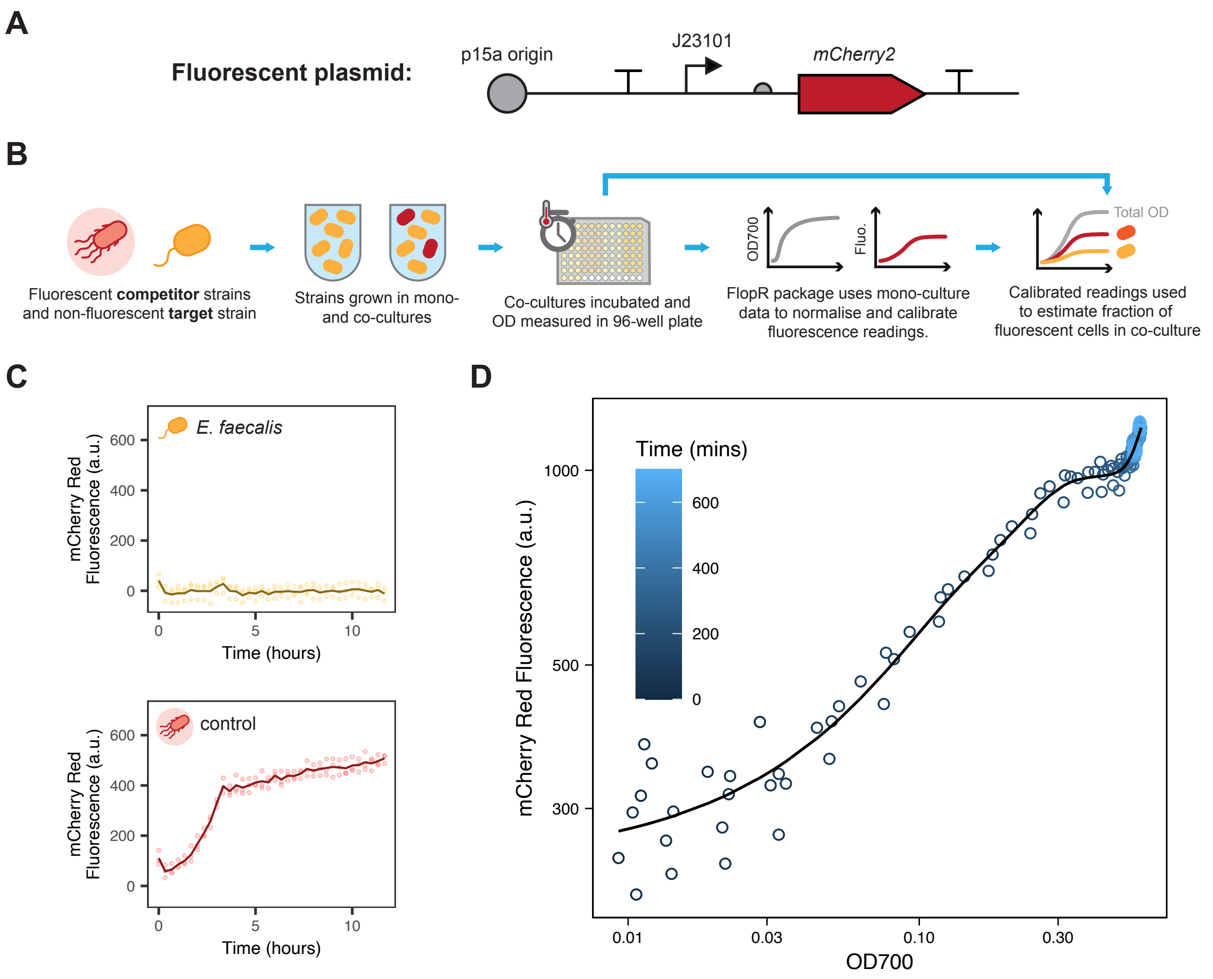

Figure S3: Summary of the FlopR process used to estimate individual species OD in co-cultures. (**A**) Layout of the mCherry2 plasmid used to produce a fluorescent signal in our competitor strains. This is required to perform FlopR analysis on co-culture timecourses. (**B**) Overview of the full FlopR assay protocol. The target and competitor strains are mixed as desired and growth measured in a 96-well plate. The mono-culture of the competitor strain is used to create a calibration curve of expected fluorescent signal for a given OD. From this calibration curve, the fluorescence measured in co-culture can be used to estimate the ratio of each species present. This ratio is used to estimate a value for the OD of each species in the co-culture. (**C**) Fluorescent mCherry signal from the target *E. faecalis* and control strain over time. It can be seen that the *E. faecalis* produces minimal fluorescence and therefore, does not interfere with the FlopR calibration process. (**D**) A representative fluorescence vs OD700 calibration curve. This curve can be used to estimate the species OD for a competitor strain from a given fluorescent signal. For a more detailed description see Fedorec *et al* (2020)[6].

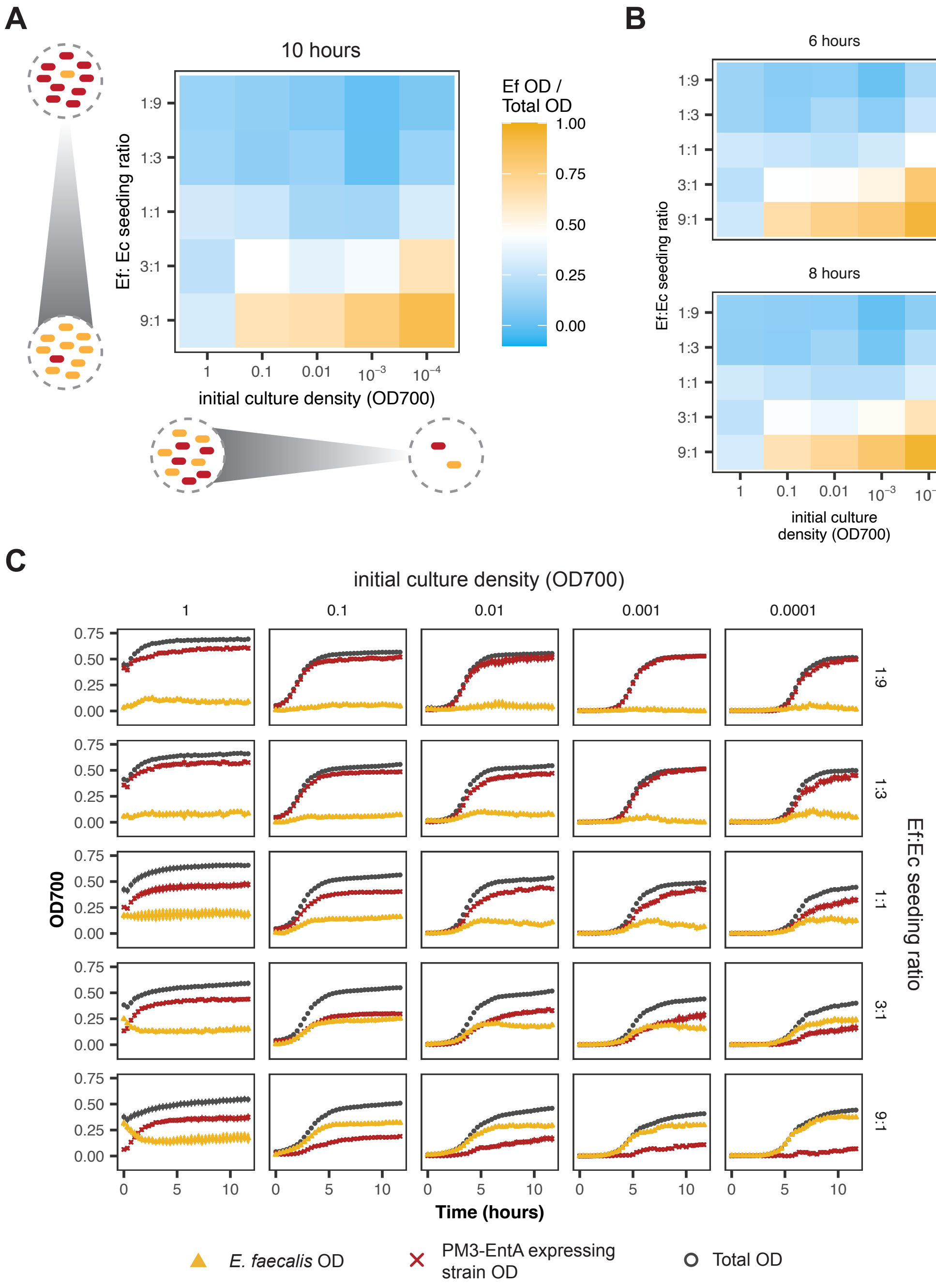

Figure S4: The effect of seeding ratio and starting density on co-culture growth. (**A**) The ten hour ratio of estimated *E. faecalis* OD over total OD for the given starting culture densities and seeding ratios. (**B**) The same estimated ratios for the 6 hour and 8 hour timepoints. (**C**) The growth curves of co-cultures from each starting condition, orange and red lines show the estimated *E. faecalis* and competitor OD, respectively. (lines indicate mean of 4 replicates ± SE)

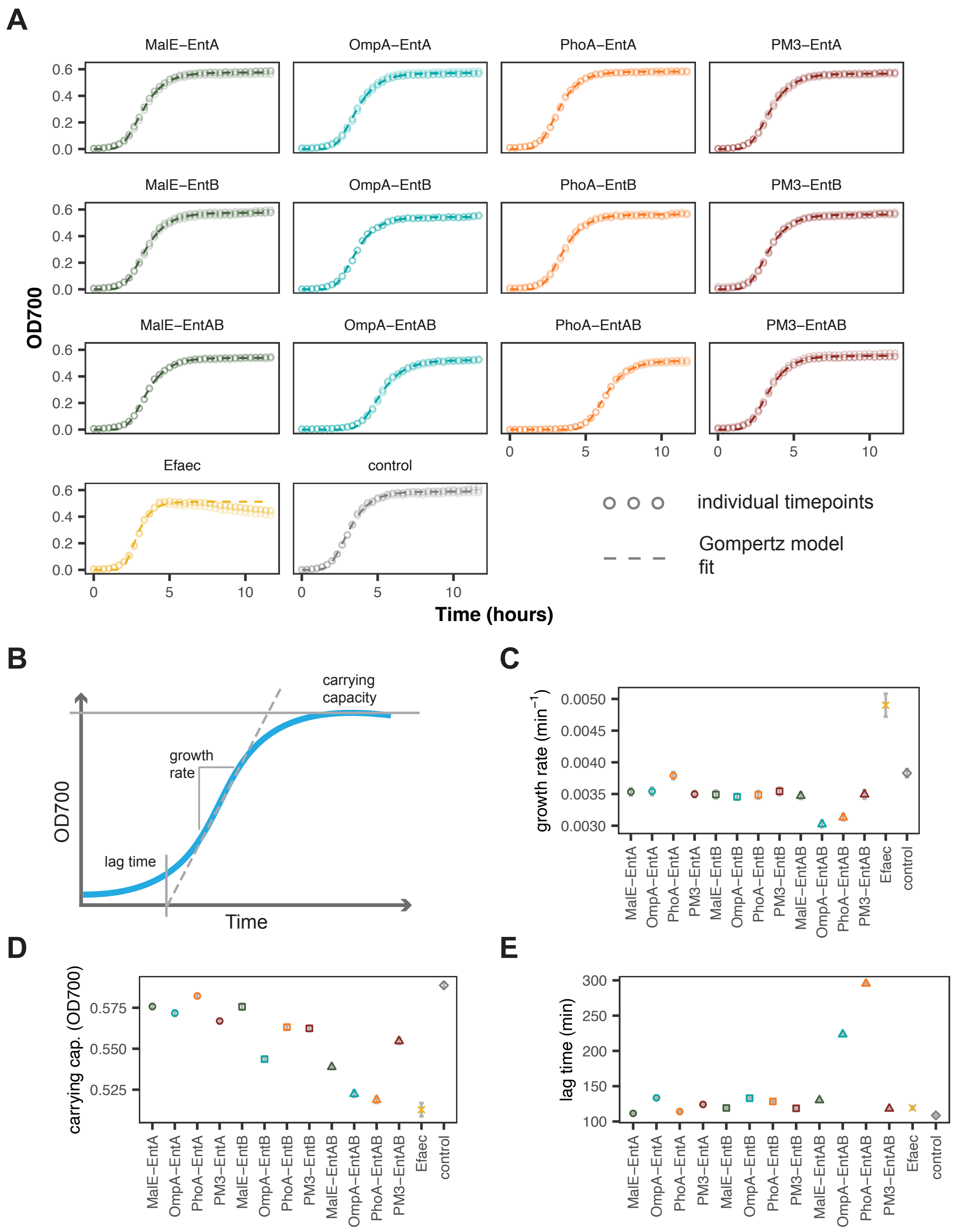

Figure S5: (**A**) Growth curves of strains grown in monoculture, fitted with the Gompertz model. Panel labels give the respective strains, 'Efaec' refers to the *E. faecalis*. Fitting of the *E. faecalis* growth curve was trimmed to 400 minutes, to exclude the effect of dropping OD700 at later timepoints (n = 3, dashed lines give model fit and points individual repeats). Parameters from Gompertz model fits of the monoculture growth curves, strains are labelled based on the bacteriocin construct they expressed: (**B**) Diagram of a typical growth curve illustrating the fitted Gompertz parameters. The fitted Gompertz parameter values for (**C**) growth rate, (**D**) carrying capacity and (**E**) lag time (fitted value ± fitted error).

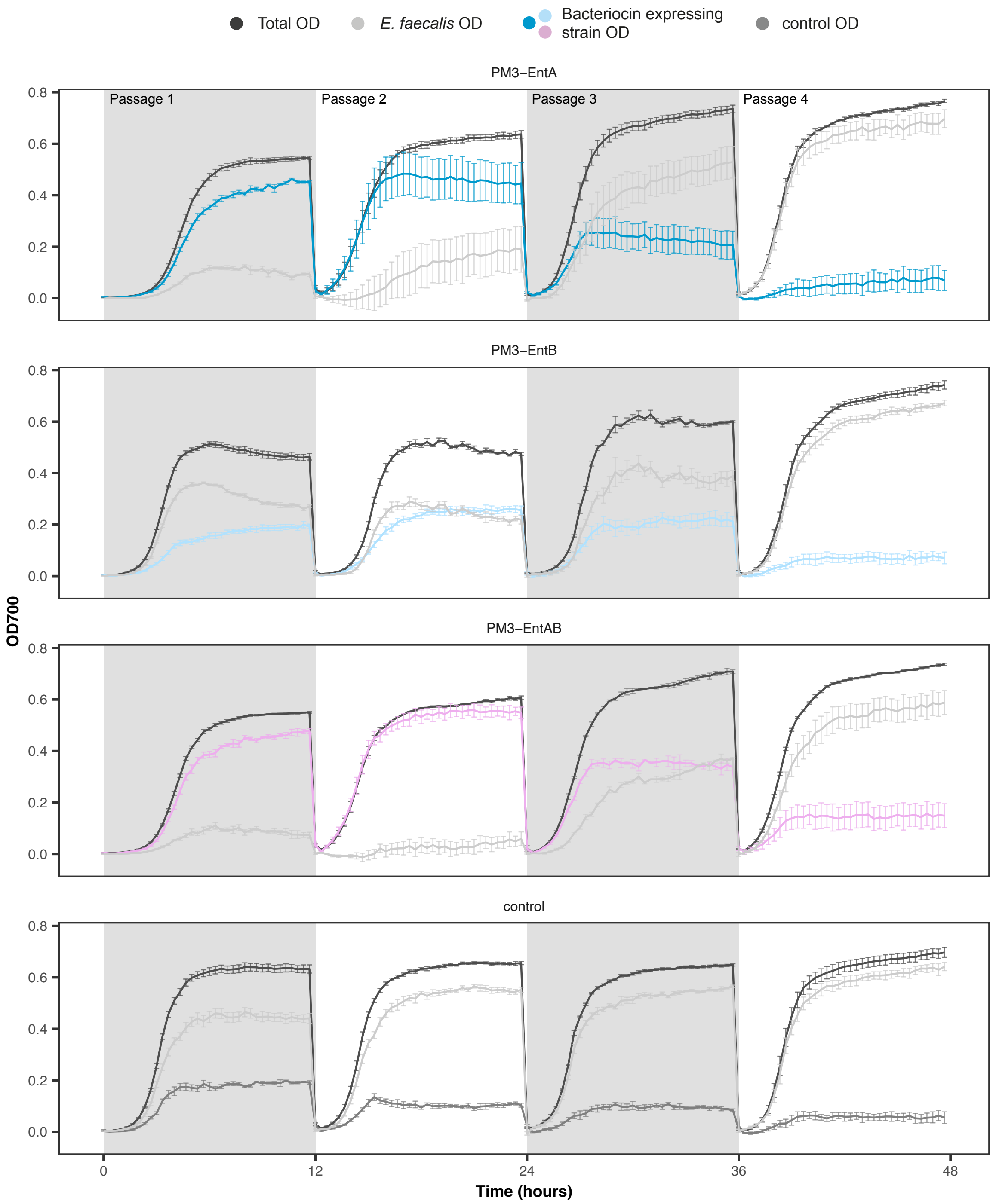

Figure S6: Co-culture time series of *E. faecalis* grown in co-culture with each of the labelled strains. Co-cultures were passaged every 12 hours, for a total of 48 hours. For all co-cultures *E. faecalis* growth increases over the 48 hour period, indicating the emergence of resistant bacteria. Anomalous datapoints were excluded from the averages, shaded areas indicate each passage (n = 4, mean values ± SE).

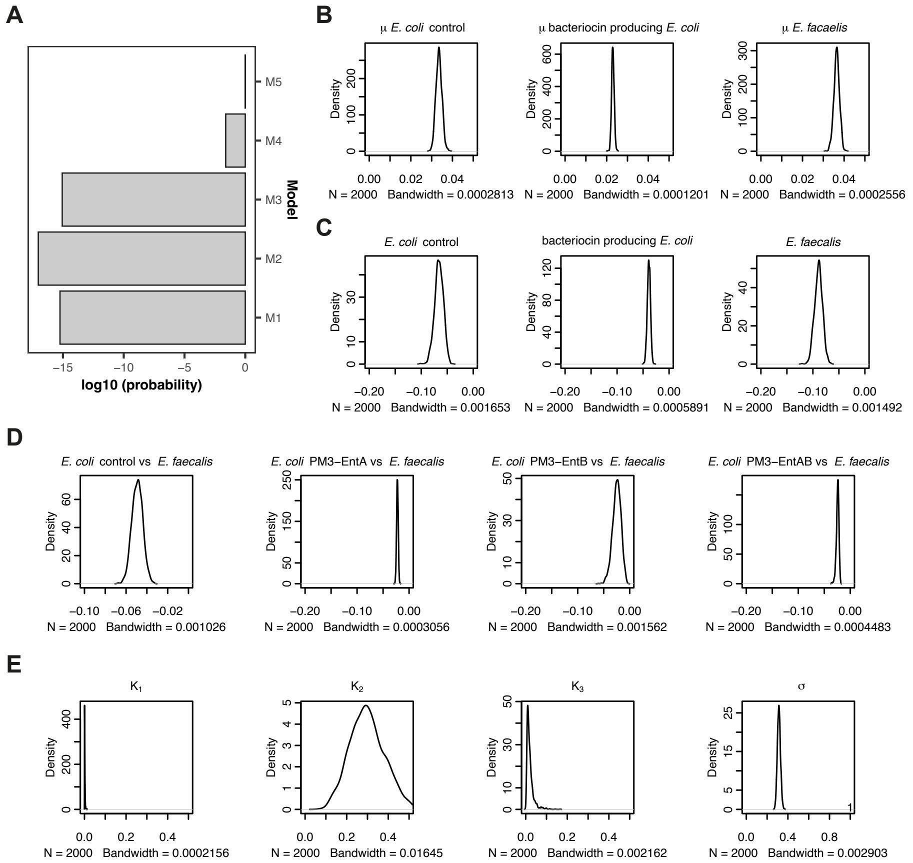

Figure S7: (**A**) The probability of each of the five models explored here. Posterior distributions for each of the parameters, calculated during Bayesian fitting of model M5: (**B**) species growth rates, (**C**) self-interaction terms, (**D**) interspecies interaction terms and (**E**) the $K_i$ and $\sigma$ parameters.

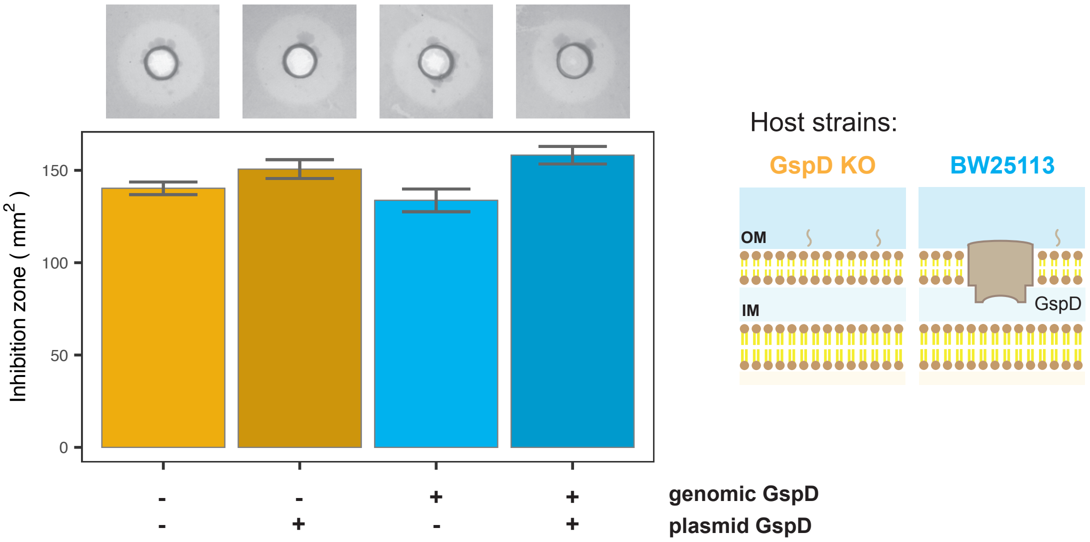

Figure S8: Inhibition assays of the PM3-EntA and PM3-EntA-GspD constructs, expressed from two host strains: the $\Delta gspD$ knockout (yellow bars), and $gspD^{+}$ BW25113 (blue bars) strains. The x-axis labels indicate plasmid and genomic expression of GspD. GspD pore deletion did not prevent bacteriocin killing under the conditions tested. Insets show representative images of the inhibition zones. (n = 3, mean values ± SE)

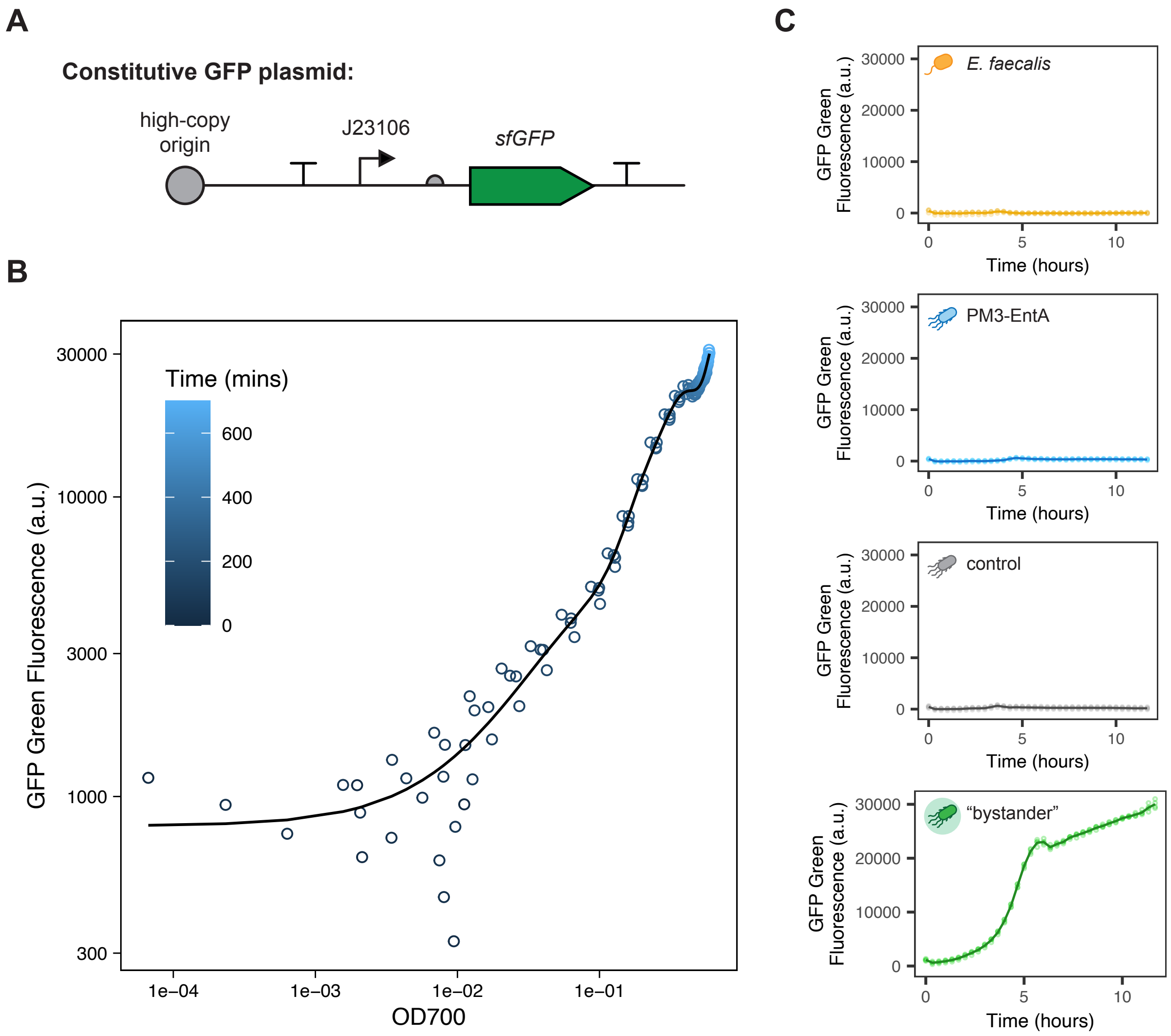

Figure S9: Summary of the incorporation of a (GFP-fluorescent) bystander strain into the liquid-culture assays. (**A**) Layout of the GFP plasmid used to produce a fluorescent signal in our bystander strain. (**B**) A representative fluorescence vs OD700 calibration curve, used to estimate the species OD of the bystander strain. (**C**) Fluorescent GFP signal from the *E. faecalis*, PM3-EntA, control and bystander strains over time. Only the bystander strain was found to produce a strong GFP signal.

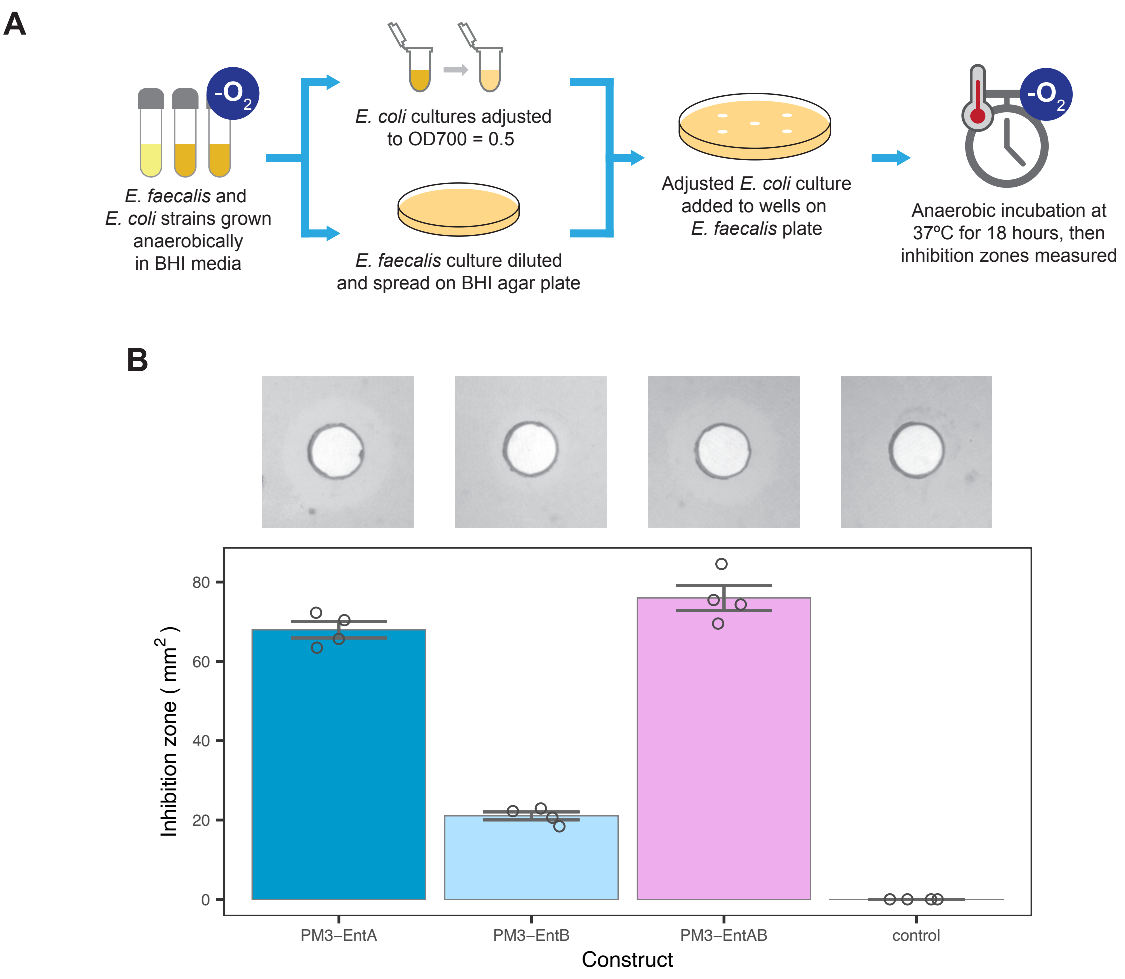

Figure S10: Inhibition zones of the PM3-bacteriocin expressing strains, under anaerobic conditions. (**A**) Overview of the anaerobic assay protocol. All cultures were grown in the absence of oxygen. (**B**) Inhibition zone assay of the PM3-bacteriocin constructs under anaerobic conditions. Inhibition zones were seen for all bacteriocin producing strains tested. Insets show representative images of the inhibition zones observed (n = 4, bars indicate mean ± SE).

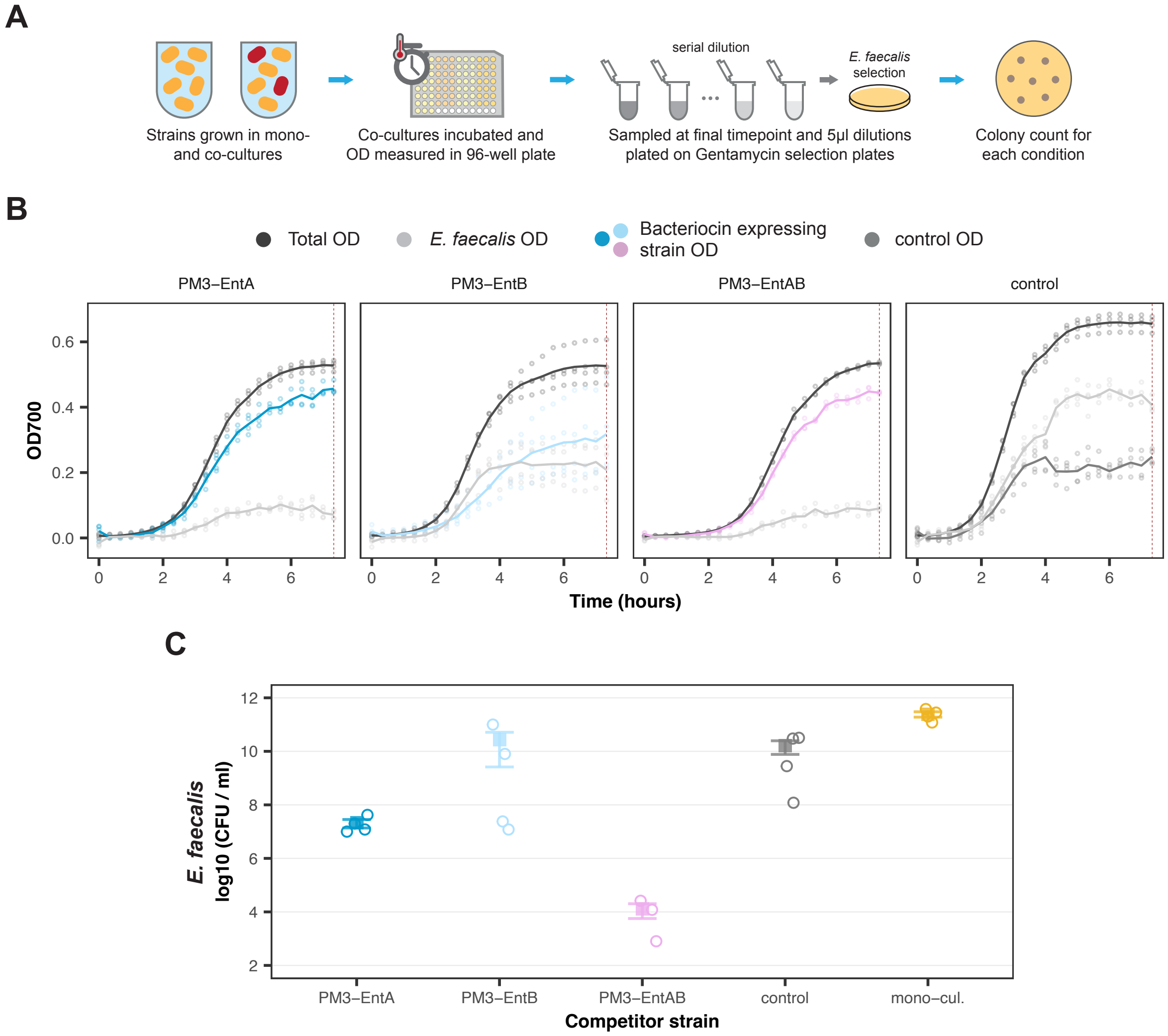

Figure S11: Colony counts of *E. faecalis* growth from FlopR co-culture assays. (**A**) The experimental protocol used for collecting estimates of colony counts from the FlopR co-culture assays. In brief, strains were prepared and grown in co-culture as described in Figure 3A. After 8 hours of growth, all co-cultures were sampled and serial dilutions plated on gentamycin containing media to select for *E. faecalis* growth. (**B**) Timecourses of the FlopR co-cultures sampled for colony counting, the red dashed line indicates the time at which samples were taken for plating ($n \geq 3$, solid lines give mean values). (**C**) The estimated colony counts of *E. faecalis* for each of the given culture conditions ($n \geq 3$, mean values $\pm$ SE).

### References

[1] Tomoya Baba, Takeshi Ara, Miki Hasegawa, Yuki Takai, Yoshiko Okumura, Miki Baba, Kirill A Datsenko, Masaru Tomita, Barry L Wanner, and Hirotada Mori. Construction of ¡i¿escherichia coli¡/i¿ k12 inframe, singlegene knockout mutants: the keio collection. *Molecular Systems Biology*, 2, 1 2006.

[2] Babak Momeni, Li Xie, and Wenying Shou. Lotka-volterra pairwise modeling fails to capture diverse pairwise microbial interactions. *Elife*, 6:e25051, 2017.

[3] Stan Development Team. RStan: the R interface to Stan, 2023. R package version 2.21.8.

[4] Bob Carpenter, Andrew Gelman, Matthew D Hoffman, Daniel Lee, Ben Goodrich, Michael Betancourt, Marcus Brubaker, Jiqiang Guo, Peter Li, and Allen Riddell. Stan: A probabilistic programming language. *Journal of statistical software*, 76(1), 2017.

[5] Felix Teufel, Jos Juan Almagro Armenteros, Alexander Rosenberg Johansen, Magns Halldr Gslason, Silas Irby Pihl, Konstantinos D. Tsirigos, Ole Winther, Sren Brunak, Gunnar von Heijne, and Henrik Nielsen. Signalp 6.0 predicts all five types of signal peptides using protein language models. *Nature Biotechnology*, 40:1023–1025, 7 2022.

[6] Alex J. H. Fedorec, Clare M. Robinson, Ke Yan Wen, and Chris P. Barnes. Flopr: An open source software package for calibration and normalization of plate reader and flow cytometry data. *ACS Synthetic Biology*, 9:2258–2266, 9 2020.
